# Key substitutions in the spike protein of SARS-CoV-2 variants can predict resistance to monoclonal antibodies, but other substitutions can modify the effects

**DOI:** 10.1101/2021.07.16.452748

**Authors:** Sabrina Lusvarghi, Wei Wang, Rachel Herrup, Sabari Nath Neerukonda, Russell Vassell, Lisa Bentley, Ann E. Eakin, Karl J. Erlandson, Carol D. Weiss

**Author notes:** These authors contributed equally to this work (Sabrina Lusvarghi, Wei Wang, Rachel Herrup, Sabari Nath Neerukonda, Russell Vassell, Lisa Bentley).

## Abstract

Mutations in the spike protein of SARS-CoV-2 variants can compromise the effectiveness of therapeutic antibodies. Most clinical-stage therapeutic antibodies target the spike receptor binding domain (RBD), but variants often have multiple mutations in several spike regions. To help predict antibody potency against emerging variants, we evaluated 25 clinical-stage therapeutic antibodies for neutralization activity against 60 pseudoviruses bearing spikes with single or multiple substitutions in several spike domains, including the full set of substitutions in B.1.1.7 (Alpha), B.1.351 (Beta), P.1 (Gamma), B.1.429 (Epsilon), B.1.526 (Iota), A.23.1 and R.1 variants. We found that 14 of 15 single antibodies were vulnerable to at least one RBD substitution, but most combination and polyclonal therapeutic antibodies remained potent. Key substitutions in variants with multiple spike substitutions predicted resistance, but the degree of resistance could be modified in unpredictable ways by other spike substitutions that may reside outside of the RBD. These findings highlight the importance of assessing antibody potency in the context of all substitutions in a variant and show that epistatic interactions in spike can modify virus susceptibility to therapeutic antibodies.

**Importance:** Therapeutic antibodies are effective in preventing severe disease from SARS-CoV-2 infection (COVID-19), but their effectiveness may be reduced by virus variants with mutations affecting the spike protein. To help predict resistance to therapeutic antibodies in emerging variants, we profiled resistance patterns of 25 antibody products in late stages of clinical development against a large panel of variants that include single and multiple substitutions found in the spike protein. We found that the presence of a key substitution in variants with multiple spike substitutions can predict resistance against a variant, but that other substitutions can affect the degree of resistance in unpredictable ways. These finding highlight complex interactions among substitutions in the spike protein affecting virus neutralization and potentially virus entry into cells.

## Introduction

Efforts to control the spread of the severe acute respiratory syndrome coronavirus 2 (SARS-CoV-2) and prevent severe SARS-CoV-2 disease (COVID-19) has led to the development of antibody-based medical countermeasures, including vaccines, convalescent plasma, and therapeutic monoclonal antibodies. Antibody therapeutics have been effective in preventing severe COVID-19 (1–3). Five neutralizing antibody products (nAbs) are in clinical use under an FDA Emergency Use Authorization (EUA), including two single nAb products (one of which the EUA has since been rescinded) (4, 5), two combination nAb products (cnAbs) (6, 7), and convalescent plasma (8). Many others are in advanced stages of clinical development (9). These products target the SARS-CoV-2 spike protein (Spike). Spike consists of a surface subunit (S1) and transmembrane subunit (S2). S1 contains an N-terminal domain (NTD) and a receptor binding domain (RBD) that mediates virus attachment to the angiotensin converting enzyme (ACE2) receptor on host cells (10–12) (Fig 1A). After S1 binds to ACE2, a cellular protease cleaves S2 to activate fusion between virus and host cell membranes, allowing the virus to enter cells (13). All mutations of concern (MOCs) identified in EUA Fact sheets as conferring resistance to nAbs are within the RBD and involve these key substitutions: P337H/L/R/T, E340A/K/G, K417E/N, D420N, N439K, K444Q, V445A, N450D, L452R, Y453F, L455F, N460K/S/T, V483A, E484K/Q/D, F486V, F490S, Q493K/R, S494P, and N501Y/T (5–7).

**Figure 1.**
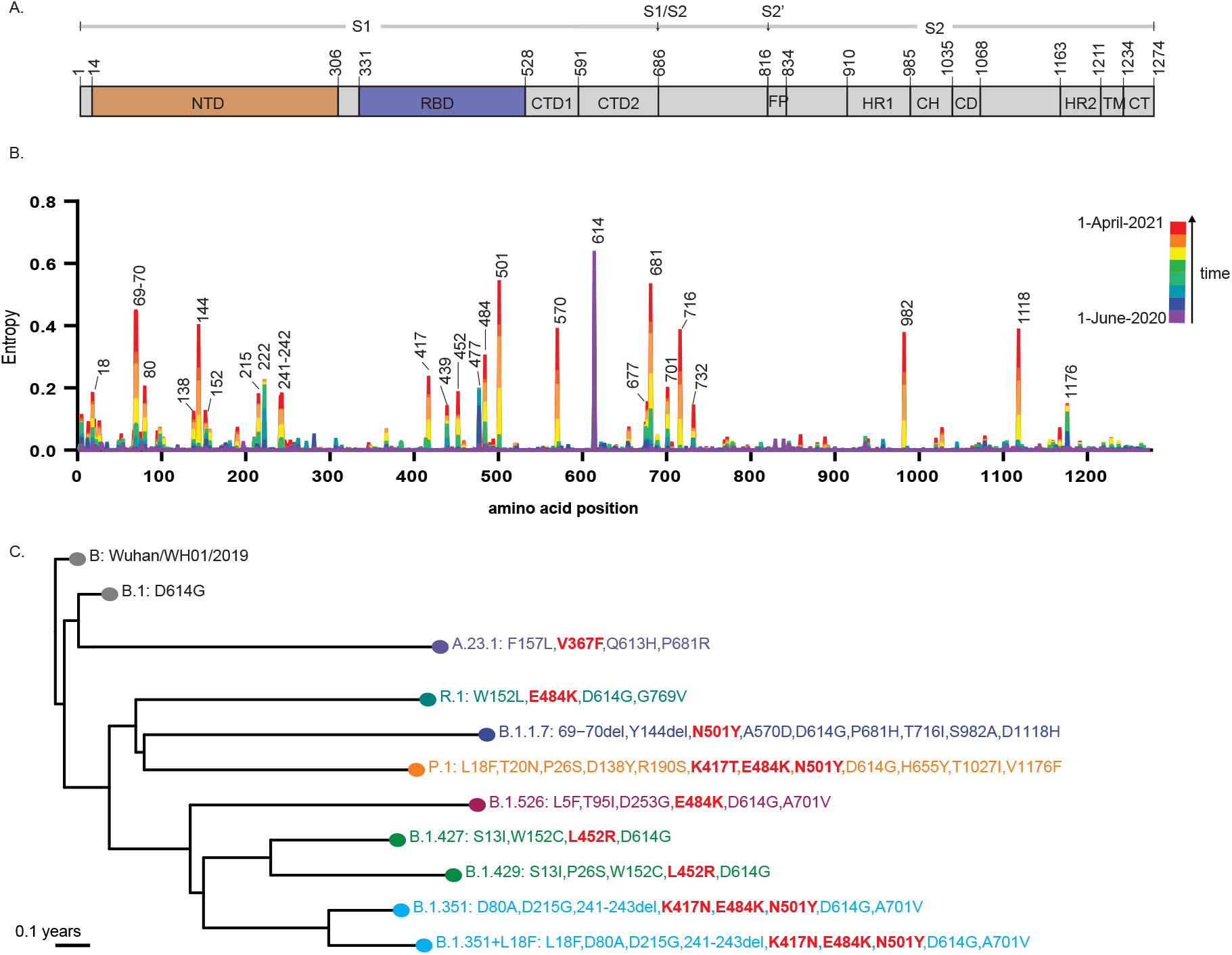
SARS-CoV-2 Spike protein domains and genetic diversity over time. (A) Schematic representation of the domains of the Spike protein precursor. SS: signal peptide; NTD: N-terminal domain; RBD: receptor binding domain; CTD1: C-terminal domain 1; CTD2: C-terminal domain 2; FP: fusion peptide; HR1: heptad repeat 1; CH: central helix; CD: connector domain; HR2, heptad repeat 2; TM, transmembrane domain; CT: central helix. Arrows indicate cleavage sites for furin (S1/S2) and TMRPSS-1 (S2’). (B) Changes in the genetic diversity of the Spike with time. Values were obtained from the normalized Shannon entropy per codon reported in GISAID at different time points. The progression of specific mutations over time in different regions of the Spike can be observed. (C). The rooted phylogenetic tree of the variants of concern (VOCs), variants of interested (VOIs), or other variants included in this study. The full genomic tree was adapted from from nextstrain/ncov (69, 70). Substitutions in the RBD are highlighted in bold red letters.

Many current nAb products were developed using SARS-CoV-2 sequences identified at the start of the pandemic (14–22). Subsequent viral evolution has led to the emergence of distinct viral lineages (23) (Fig 1B and C and Table S1), raising concerns about the continued effectiveness of many nAbs. One of the earliest variants that became globally dominant contains the D614G Spike substitution (24, 25). Subsequently, the B.1.1.7 (also known as Alpha), B.1.351 (also known as Beta), P.1 (also known as Gamma) and A.23.1 lineages evolved independently in the United Kingdom, South Africa, Brazil, and Uganda, respectively. Additional variants of concern (VOC) or variants of interest (VOI) have emerged regionally in the United States (US), including B.1.429/B.1.427 variants (also known as Epsilon) first identified in California and B.1.526 variants (also known as Iota) first identified in New York (26).

B.1.1.7 lineage variants are highly contagious (27, 28) and have become dominant throughout the US (23). Although B.1.1.7 variants contain the N501Y substitution in the RBD and three deletions in the NTD, studies suggest that COVID-19 vaccinee sera and some monoclonal antibodies have only slightly reduced potency against this variant (29–31). Antibody products available under EUA have no change in susceptibility against B.1.1.7. Novel variants of the B.1.1.7 lineage have acquired additional MOCs, including E484K and S494P substitutions. The B.1.427/B.1.429 variants have three substitutions in Spike, including one MOC involving the L452R substitution. Vaccinee sera effectively neutralized B.1.427/B.1.429 variants with similar potency against B.1.1.7, but the L452R substitution abolished potency of some nAbs (32, 33).

The B.1.351 lineage variants have three RBD MOCs involving K417N, E484K, and N501Y substitutions, and five non-RBD mutations along with a deletion in the NTD. The P.1 variant contains very similar RBD MOCs involving K417T, E484K, and N501Y substitutions, along with nine mutations in non-RBD regions. Several studies have reported a significant drop in the neutralization potency of convalescent and vaccinee sera, as well as monoclonal antibodies, against B.1.351 and P.1 variants (33–41). B.1.526 variants contain E484K and five mutations in the non-RBD region. Convalescent and vaccinee sera, as well as at least one therapeutic nAb, displayed a significant drop in the neutralization activity against B.1.526 variants, though a cocktail of nAbs retained neutralization potency (42, 43). R.1 lineage variants with E484K and non-RBD amino acid substitutions appeared in Japan and USA, and the B.1.617.1 and B.1.617.2 variants (also known as Delta) recently identified in India and spreading worldwide contains the E484Q and S478K mutations, respectively.

The ongoing evolution of SARS-CoV-2 requires frequent assessments nAb potency against variants. As it can take weeks to acquire new Spike genes and viruses for performing *in vitro* evaluations, information about substitutions that can predict resistance to nAbs is needed. This study was a US government effort to profile nAbs in late stages of clinical development for resistance against a large panel of pseudoviruses representing SARS-CoV-2 variants with single and multiple mutations. Findings from this study about the effects of these substitutions inform the clinical use and development of new nAbs and provide new insights into substitutions that can affect antibody binding to Spike and potentially virus entry into cells.

## Results

### Choice of spike mutations and therapeutic antibodies for study

This comparative profiling study was initiated in October 2020 by the US government COVID-19 Response Therapeutics Research Team to guide the clinical testing and use of nAbs. Twenty-five nAbs in late stages of clinical investigation were donated by developers with an agreement that data would be kept blinded except to the study investigators and those who signed a confidentiality agreement. Participating developers were provided unblinded data only for their own products and a blinded heatmap showing the fold-change compared to wildtype for all products. Due to the agreement terms, nAbs described here are only shown with randomized identification codes as follows: single nAbs (nAbs.A to R), combination of two nAbs (cnAbs.S to X), and polyclonal nAbs (pnAbs.I to IV).

The nAbs studied were developed early in the pandemic based on sequences from the first SARS-CoV-2 viruses available. Spike substitutions chosen for study were initially based on early variants reported in the Global Initiative on Sharing All Influenza Data (GISAID) database (Fig S1) or experimental studies (44–50). Our goal was to set up a standardized assay and set of mutations to look at the entire portfolio of nAb products as a matrix to proactively and methodically identify Spike substitutions that had potential to escape neutralization across the nAbs. This approach allows identification of gaps and duplications in coverage across the portfolio, as well as potential novel nAb combinations. In the past year, multiple mutations, especially involving the RBD, NTD, and the furin cleavage site, have accumulated as the virus has evolved into distinct lineages (Fig 1B and C and Table S1).

### Resistance patterns against pseudoviruses with single amino acid substitutions in Spike

Using the D614G as the wildtype Spike (WT), we first screened nAbs against pseudoviruses bearing Spikes with single amino acid substitutions or NTD deletions to identify key substitutions that could confer resistance to any nAb (Fig 2 and Fig S2). While our assay precision allowed us to reliably detect three-fold differences (51), we used 10-50-fold and >50-fold increase in the IC_50_ relative to WT as screening benchmarks for modest and complete loss of activity, respectively (Fig 2A). We chose these cutoffs because therapeutic levels of monoclonal antibodies may be high enough to overcome low levels of resistance. Nonetheless, the clinical relevance of the changes in IC_50_ has not been determined for any *in vitro* assay, and the clinical efficacy for most of the nAbs tested has not been established. The ratios do not account for the absolute potency of a given nAb. Several nAbs failed to neutralize some variants at the highest concentrations tested (Fig 2B), suggesting that a protective antibody concentration may not be physiologically achievable for that variant. Figure S2 provides the fold IC_50_ change relative to WT for all nAbs.

**Figure 2.**
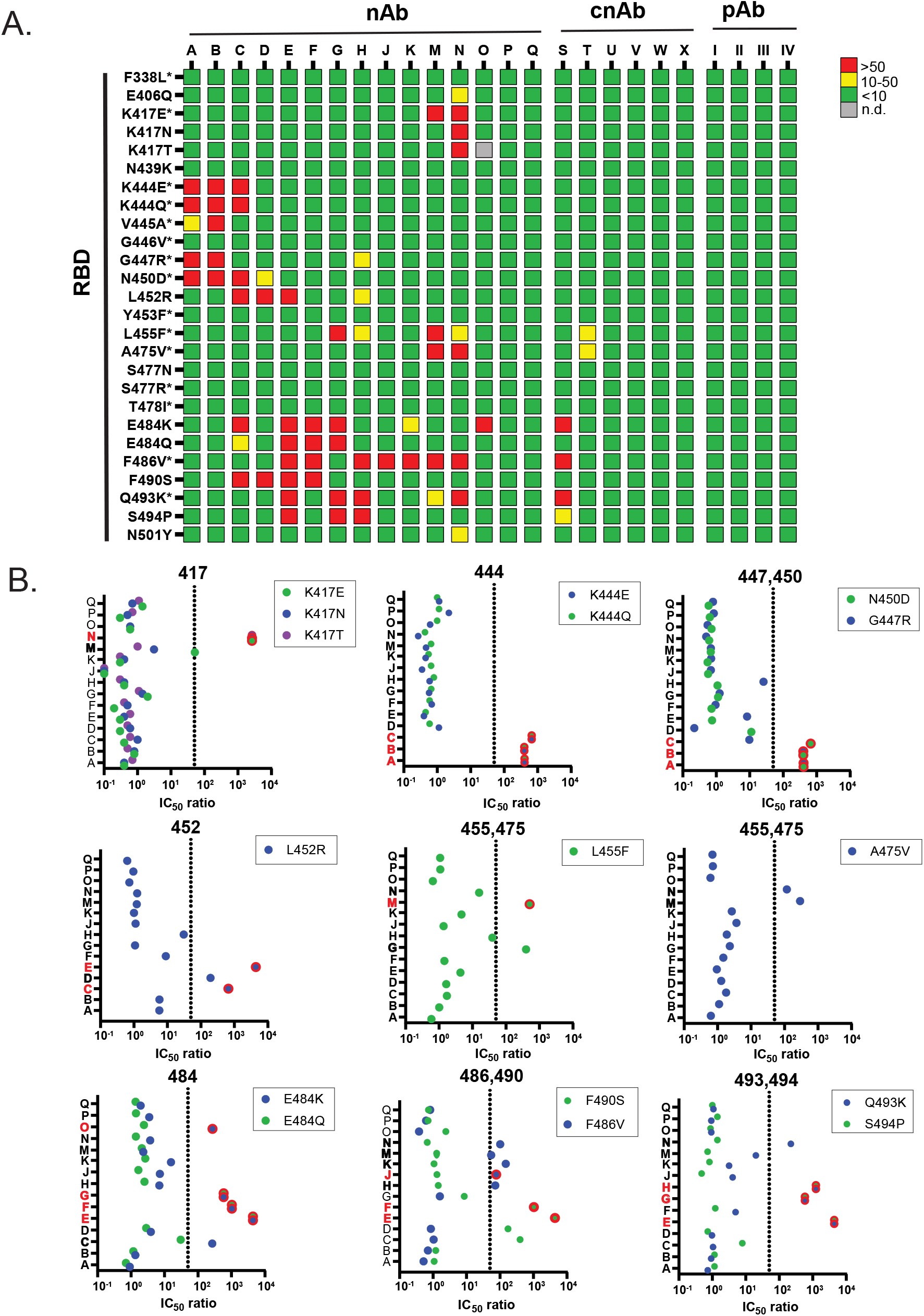
Resistance patterns of therapeutic antibodies conferred by single substitutions in the RBD of the SARS-CoV-2 spike protein. The blinded antibody panel, consisting of 15 single nAbs, six cnAbs, and four pAbs were tested against 26 pseudoviruses with the indicated single substitutions in the D614G Spike. (A) Heat map representing the ratio of IC_50_ values of the variant relative to wild type. Red indicates loss of potency (IC_50_ ratios > 50). Yellow indicates moderate loss of potency (IC50 ratios between 10-50), and green indicates retention of potency (IC_50_ ratios <10). The gray square indicates not done (n.d.). The asterisk (*) indicates mutations not reported to GISAID. (B) Dot plots show IC_50_ ratios of individual substitutions that have a significant impact on one or more antibodies. Dots outlined with red circles indicate that the highest concentration tested was not sufficient to achieve full neutralization of the pseudovirus with the indicated substitution in Spike. Red letters highlight the nAbs that did not neutralize the indicated pseudoviruses at the highest concentration tested. Bold letters highlight the nAbs that had at least a 50-fold reduction in potency (shown by dotted line) to the indicated pseudoviruses. Data shown represent at least two independent experiments each with an intra-assay duplicate.

We evaluated 48 pseudoviruses with single substitutions or deletions in Spike (Fig S2), including 20 in the NTD, 26 in the RBD, and two in more C-terminal regions of Spike. Single substitutions conferring resistance resided exclusively in the RBD (Fig 2A, top panel), reflecting the immunodominance of this region and the way most nAbs and cnAbs were selected. The tested NTD changes did not significantly change the potency of any nAb. Against the 26 pseudoviruses with single RBD substitutions, only six substitutions (F338L, N439K, G446V, Y453F, S477N/R and T478I) did not affect the potency of any nAb. Only two (nAbs.P and Q) of the 15 nAbs retained potency (<10-fold change compared to WT) against all pseudoviruses with single Spike substitutions.

Single substitutions residing in positions 484-494 and 444-452 of the RBD conferred resistance to 14 of the 15 single nAbs (Fig 2). The residues in these regions surround, and in some cases overlap, the ACE2 binding sites and affect many nAbs (Fig 3). Several substitutions were based on *in vitro* experiments (44–50) that have not yet been reported in natural variants. These pseudoviruses were functional (Fig S3), indicating that they have the potential to emerge and impact some nAbs. Consistent with many reports (38, 52–60), the E484K substitution present in many lineages had a major impact on many nAbs, leading to a loss of potency for five nAbs (nAbs.C, E, F, G, and O) and a modest loss of potency for nAb.K (Fig 2). The E484Q substitution, present in B.1.617.1 and B.1.617.3 variants, attenuated the potency of four nAbs that were affected by E484K. Interestingly, nAb.O retained neutralization against E484Q but not E484K.

**Figure 3.**
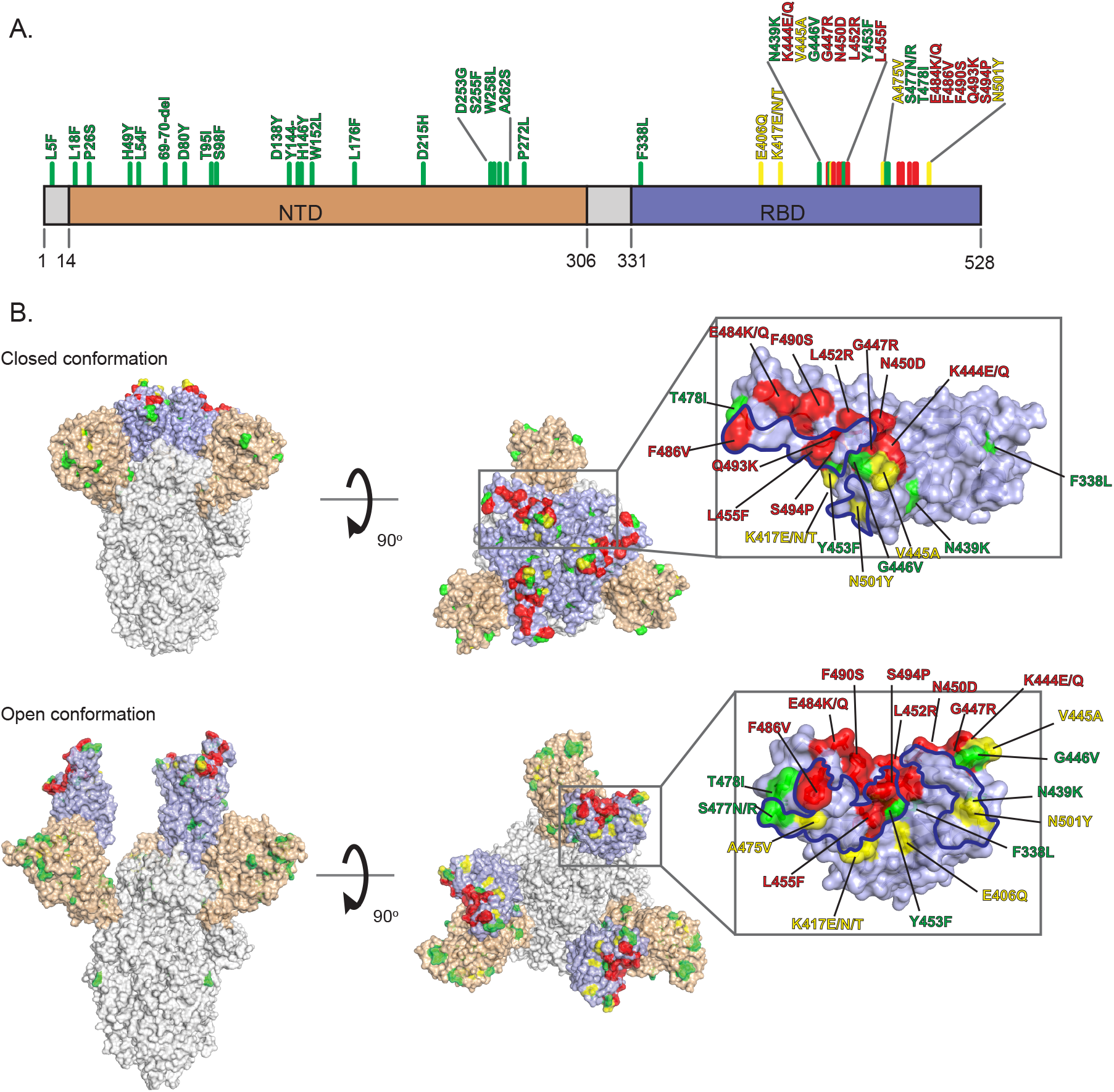
Hot spots for substitutions in the receptor binding domain (RBD) that confer resistance to therapeutic antibodies. (A) Spike domains NTD (light orange) and RBD (light purple) are shown with the amino acid substitutions color-coded according to the number of nAbs affected. (B) Spike 3D structures shown in the closed (PDBID: 6ZOZ) and open (PDBID: 7A98) (side view and top view) conformations. Green residues are the substitutions that do not affect the potency of any nAbs tested. Yellow residues are the substitutions that reduce the potency of one-to-two nAbs by at least 10-fold. Red residues are the substitutions that reduce the potency of three or more nAbs by at least 10-fold. The RBD region are zoomed in the right insets. The amino acids in the RBD region that within 4Å of ACE2 binding sites are delineated with a blue line.

The largest number of nAbs were affected by the F486V substitution, which conferred a loss of potency for eight nAbs (nAbs.E, F, H, J, K, L, M, and N). This substitution was previously identified in an *in vitro* selection (45) but has not yet been reported in GISAID. Other substitutions that reduced potency were: F490S, which conferred a loss of potency for four nAbs (nAbs.C, D, E, and F); Q493K, which conferred a loss of potency for four nAbs (nAbs.E, G, H, and N) and a modest reduction of potency for nAb.M; and S494P, which conferred a loss of potency for three nAbs (nAbs.E, G, and H). The F490S and S494P substitutions have been found in different lineages.

Single substitutions in residues 444-452 located on a different side of the receptor binding motif (Fig 3) also reduced the potency of many nAbs, but only substitutions involving L452 have been consistently found in several lineages, namely the B.1.427, B.1.429, and B.1.617 variants. The L452R and N450D substitutions each conferred a loss of potency for three nAbs (nAbs.C, D, and E and nAbs.A, B, and C, respectively) and a modest loss of potency for another (nAbs.H and D, respectively). The K444E/Q substitutions each conferred a loss of potency for the same three nAbs (nAbs.A, B, and C). Other substitutions affecting two or more nAbs were: A475V, which conferred a loss of potency for two nAbs (nAbs.M and N); L455F, which reduced potency of four nAbs (nAbs.G (loss), H, M (loss), and N); and V445A, which reduced potency of nAb.A and conferred a loss of potency for nAb.B. Among the tested substitutions between residues 475-478, only A475V conferred a loss of potency for two nAbs (nAbs.M and N). Mutations affecting residues 477 and 478 have been found in different lineages, but they did not affect the potency of the nAbs.

The substitutions at residue 417 conferred resistance or enhanced sensitivity to neutralization for a few nAbs (Fig 2B and Fig S1). NAb.N was resistant to K417E/N/T, and nAb.M was resistant only to K417E. By contrast, nAb.J was more potent against K417E/N/T and T478I, and nAb.F was more potent against K417E. The dual effect of the K417N substitution for various antibodies was previously seen by others (38, 54).

Several nAbs that lost potency against pseudoviruses with Spike substitutions were not able to neutralize variants at the highest concentrations tested (Fig. 2B, dots with red outline). Resistance at high concentrations of nAbs (> 50-fold increase relative to WT) may predict a loss of potency *in vivo*. This occurred for: nAbs.M and N for substitutions for K417E and nAb.N for K417N/T; nAbs.A, B, and C for K444E/Q; nAbs.A, B, and C for N450D; nAbs.C, D, and E for L452; nAb.G and M for L455F; nAb.M and N for A475V; nAbs.C, E, F, G, and O for E484K; nAbs.E, F, and G for E484Q; nAbs.E, F, H, J, K, L, M, and N for F486V; nAbs.C, D, E, and F for F490S; nAbs.E, G, H and N for Q493K; and nAbs. E, G, and H for S494P.

Importantly, all pnAbs and four of six cnAbs (cnAbs.U, V, W, and X) retained potency against the pseudoviruses with the RBD mutations. CnAb.S lost potency against pseudoviruses with the E484K, F486V and Q493K substitutions and had reduced potency against the pseudoviruses with the S494P substitution (Fig 2A and Fig S2). CnAb.T had modestly reduced potency due to L455F and A475V substitutions. The cnAb that did not retain potency was comprised of two nAbs that shared susceptibility to a few substitutions, while the cnAbs that retained full potency were comprised of two nAbs with complementary resistance profiles. These resistance profiles provide new information for identifying novel complementary nAb combinations that may be resilient against the substitutions that we studied.

Altogether, the majority of the nAbs tested in this study had reduced potency against pseudoviruses with single amino acid substitutions in the RBD, but most cnAbs and all pnAbs remained potent. Many nAbs were affected by the same substitutions (Fig 3), indicating shared hotspots for nAb binding in these immunodominant regions. This analysis underscores the importance of using cocktails of nAbs with complementary resistance profiles.

### Resistance patterns against pseudoviruses representing variants of concern or interest

We next profiled the potency of the nAbs against pseudoviruses representing 11 circulating VOCs, VOIs, or other variants with mutations in the RBD (Fig 1C, and Table S1). Overall, nine nAbs lost potency against at least one VOC or VOI, including seven that lost potency to at least six variants (Fig 4A). All nAbs retained potency against the A.23.1+D614G variant. A.23.1 has the single V367F substitution in the RBD. All other variants tested contain different combinations of RBD substitutions K417N/T, L452R, E484K, S494P, or N501Y. V367F is more distant from the receptor binding motif (RBM) compared to the other RBD substitutions.

**Figure 4.**
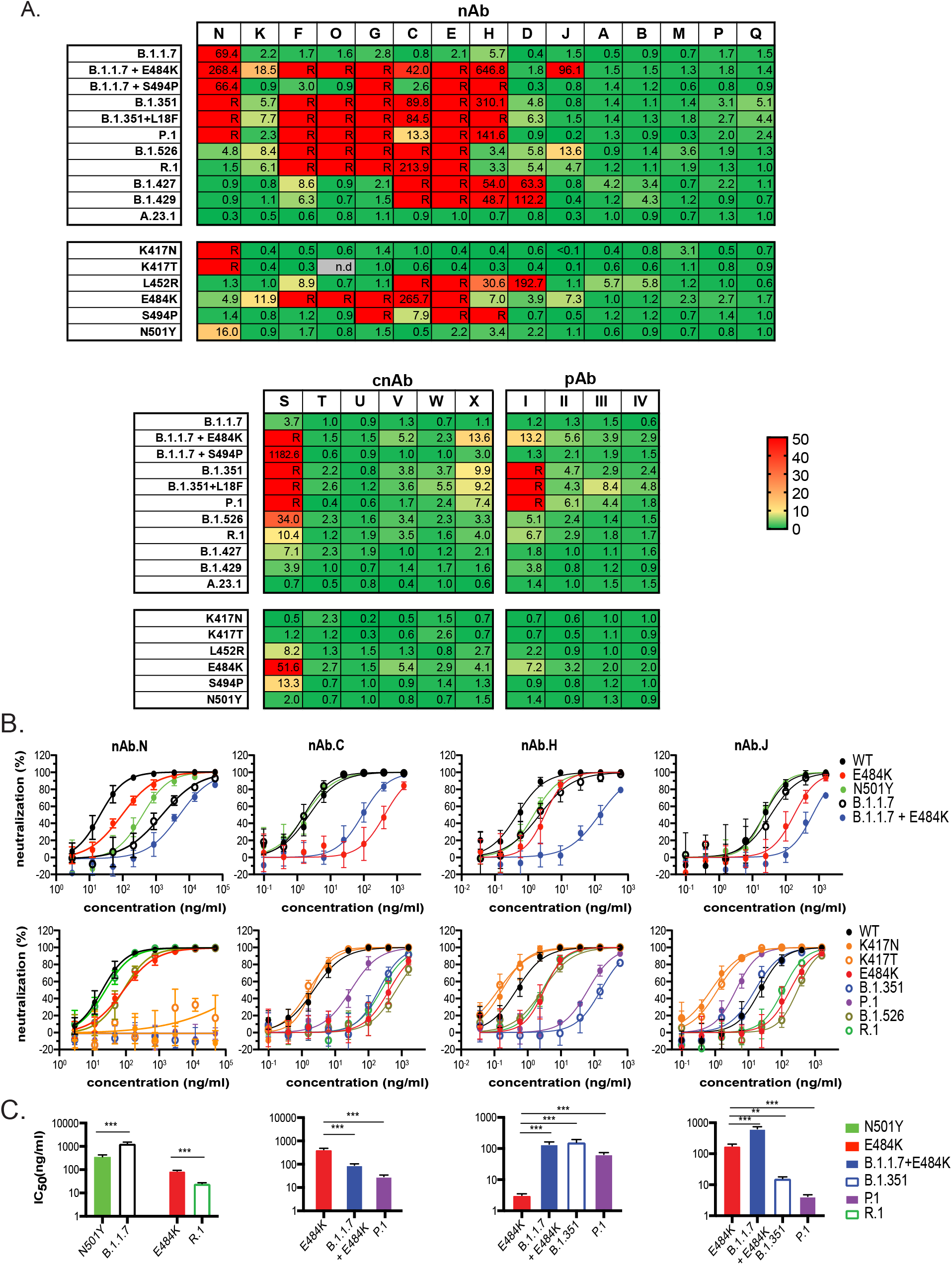
Resistance conferred by key substitutions can be modified in the context of variants with other spike substitutions. The blinded antibody panel consists of 15 nAbs, 6 cbAb and 4 pAb. (A) Heat map with the ratio between the IC_50_ of each variant with multiple substitutions or a single substitution and the IC_50_ of WT. Shades of green to yellow, yellow to orange, and red indicate ratios <10, 10-50 and >50, respectively. The IC_50_ ratios that could not be determined due to incomplete neutralization at the highest concentration tested are listed as resistant (R). (B) Neutralization curves or (C) IC_50_ bar graphs of antibody-virus pairs highlighting examples where the potency of the nAb against the pseudovirus variant could not be fully predicted by a key substitution (mutation of concern) alone. The potency of the nAb in these examples was modified by the context of the key substitution with other Spike substitutions. Spikes sequences of variants with RBD substitutions highlighted in bold are listed below: B.1.1.7: 69-70deletion, 144deletion, **N501Y,** A570D, D614G, P681H, T716I, S982A, D1118H B.1.351: D80A, D215G, Del241-243, **K417N**, **E484K**, **N501Y**, D614G, A701V P.1: L18F, T20N, P26S, D138Y, R190S, **K417T**, **E484K**, **N501Y**, D614G, H655Y, T1027I, V1176F B.1.526: L5F, T95I, D253G, **E484K**, D614G, A701V R.1: W152L, **E484K**, D614G, G769V B.1.427: S13I, W152C, **L452R**, D614G B.1.429: S13I, P26S, W152C, **L452R**, D614G A.23.1: W152L, **E484K**, D614G, G769V Data shown represent at least two independent experiments each with an intra-assay duplicate. * p < 0.05, ** p < 0.01, *** p < 0.001.

All nAbs retained potency against B.1.1.7 (NTD: 69-70 deletion, 144 deletion; RBD:N501Y) except nAb.N. Interestingly, the loss of potency by nAb.N against B.1.1.7 could only be partially explained by the degree of resistance conferred by N501Y alone (Fig 4B, nAb.N top panel and Fig 4C first panel). Because N501Y is the only RBD substitution in B.1.1.7 that differs from WT, the shift in nAb.N potency against B.1.1.7 points to epistatic interactions involving non-RBD substitutions that modify nAb.N interactions with the RBD. By contrast, the loss of nAb.N potency against B.1.351 and P.1 appears to be largely due to the resistance conferred by the K417N/T substitutions, with some contribution by N501Y (Fig 4A and Fig 4B, nAb.N bottom panel). Unexpectedly, resistance conferred by E484K substitution was reversed in the context of R1 (Fig 4B bottom panel and Fig 4C first panel) ) either by the addition of the G769V substitution or by the combination of both W152L and G769V, as W152L alone only had a modest effect on nAb.N (variant/WT ratio was 1.2, Fig S2).

NAb.K. retained potency against all VOCs and VOIs except when E484K was present. The moderate loss of potency against B.1.1.7+E484K was consistent with the loss of potency against E484K alone (Fig 4A). The loss of potency for nAbs.F, O, G, and E against the various VOCs and VOIs can also be mostly explained by the presence of either E484K or S494P alone. Likewise, the loss of potency for nAb.D against B.1.427/9 can be explained by the L452R substitution alone (Fig 4A).

Differences in potency of nAb.C, nAb.H, and nAb.J against the VOCs and VOIs relative to the potency against single substitutions contained in those variants also suggested epistatic effects. While L452R alone explained the loss of potency of nAb.C against B.1.427/29 (Fig 4A), VOCs and VOIs with E484K were either more or less susceptible to neutralization by nAb.C compared to E484K alone. For B.1.1.7 + E484K and P.1 variants, the potency of nAb.C was higher (6 and 13-fold, respectively) compared to E484K alone (Fig 4B top and bottom panels and Fig 4C second panel). This result indicates that other residues in these variants offset some of the resistance conferred by E484K. Moreover, nAb.C was more potent against P.1 than against B.1.351 despite each having similar RBD substitutions (Fig 4B nAb.C bottom panel and Fig 4C nAb.C second panel). Both P.1 and B.1.351 variants share N501Y and E484K substitutions that are present in B.1.1.7+E484K, but P.1 and B.1.351 also have either K417N or K417T as a third RBD substitution. Because nAb.C has similar potency against K417N or K417T substitutions alone, the differences in nAb.C potency against P.1 and B.1.351 are not necessarily explained by the individual contributions of the RBD substitutions (Fig 4A nAb.C top panel, Fig 4B nAb.C bottom panel, and Fig 4C nAb.C second panel). Rather, the findings suggest epistatic interactions within the RBD, outside the RBD, or both.

NAb.H lost potency against B.1.1.7+E484K, B.1.1.7+S494P, B.1.351 with or without 18F, and P.1, and had moderately reduced potency against B.1.427/B.1.429 (Fig 4A). The moderately reduced potency against L452R alone explained a similar loss of potency against B.1.427/B.1.429, and the complete loss of potency against S494P alone explained the loss of potency against B.1.1.7+S494P (Fig 4A). While nAb.H retained potency against K417N/T, E484K, and N501Y substitutions alone, the potency of nAb.H against variants containing these substitutions depended on the context of other substitutions in the variants that may include substitutions or deletions outside of the RBD. The potency of nAb.H against B.1.526 and R.1 was similar to the potency against E484K alone (~ 7 fold), yet the greater loss of potency (>140 fold) against B.1.1.7+E484K, B.1.351, and P.1 could not be fully accounted for by E484K alone (Fig 4B nAb.H, top and bottom panels and Fig 4C nAb.H third panel). The N501Y substitution is present in B.1.1.7, B.1.351, and P.1, but N501Y alone in the context of WT did not have a major impact on nAb.H potency (~ 3 fold, Fig S2 and Fig 4A bottom panel). Both B.1.351 and P.1 have substitutions at residue 417, but N417K and N417T alone modestly enhanced nAb.H potency (< 0.3 fold, Fig S2). Altogether these finding indicate that functional interactions involving N501Y and E484K, possibly with other residues within RBD or non-RBD regions, are affecting nAb.H potency.

The potency of nAb.J was also reduced more by E484K in the context of B.1.1.7 than in the context of WT, but this effect was not seen for the other VOCs or VOIs (Fig 4B, nAb.J top and bottom panels and Fig 4C nAb.J fourth panel). However, in the context of B.1.351 and more so in the context of P.1, nAb.J had increased potency compared to E484K alone, B.1.526, or R.1. The N501Y substitution is shared by B.1.1.7, B.1.351, and P.1, while P.1 and B.1.351 each also have the N417T and N417K substitutions, respectively. Both N417T and N417K increased the potency of nAb.J to a similar extent (Fig 4A and B nAb.J panels), which may help to offset the effect of E484K. However, the differences in nAb.J potency against P1 and B.1.351 may be due to differences in non-RBD residues.

NAbs.A, B, M, P, and Q maintained potency against all tested VOCs and VOIs (Fig 4A). However, nAbs.A, B, and M were vulnerable to several single RBD substitutions (Fig 2A and Fig S2) that have not yet been identified in a VOC.

The cnAbs and pnAbs were generally more resilient against the VOCs and VOIs compared to the nAbs. Only cnAb.S lost potency against B.1.1.7 with E484K or S494P, B.1.351 with or without L18F, and P.1. CnAb.S also had a moderate loss of potency against B.1.526 and R.1. Each of the nAbs in cnAb.S lost potency to several RBD substitutions. The combination of multiple resistance substitutions in nearby and overlapping residues likely account for the loss of cnAb.S potency against these variants. CnAb.X and pnAb.I had modestly reduced potency against B.1.1.7+E484K, and pnAb.I lost potency against B.1.351 with or without L18F, and P.1.

In summary, variants containing E484K, S494P, or L452R conferred a loss of potency for eight of the 15 nAbs, one cnAb and a single pAb, regardless of the variant. For many nAbs, these key substitutions can serve as markers of resistance in the variants, but the degree of resistance may be significantly modified by other co-existing substitutions. We also note that not all of the combination products were better than the monotherapy products; for instance, cnAb.S lost potency against many VOC/VOIs while some single nAbs (nAb.A, nAb.B, nAb.L, nAb.M, nAb.P, nAb.Q) retained potency. Nonetheless, combination products potentially offer wider coverage against potential variants and should include potent partners with different resistance profiles.

### Antigenic cartography based on therapeutic antibodies

To visualize the antigenic relatedness of the pseudoviruses with the various Spike substitutions, we performed an antigenic cartography analysis using the nAb neutralization titers. The antigenic map shows several pseudovirus clusters (Fig 5). B.1.1.7 is antigenically close to the WT, but B.1.1.7+E484K is distant from WT and closer to E484K. B.1.1.7+S494P is also distant from WT and close to S494P. Both shifts of B.1.1.7 highlight the antigenic dominance of the E484K or S494P substitutions. The dominant antigenic effect of E484K is also evident by the relatively close clustering of pseudoviruses representing P.1, B.1.526, and B.1.351 variants with the E484K pseudovirus. However, each of these variants is a little distant from each other, indicating that that the dominance of E484K is modified by other residues in those variants. The B.1.427/29 variants containing L452R are also distant from WT and cluster with L452R in a separate region from the E484K cluster. Two other clusters are formed by either N444D/Q and N450D or L455F and F486V that lie on opposite sides of the WT cluster. The residues defining these antigenic clusters are highlighted on the different regions of the RBD (Fig 5 inset).

**Figure 5.**
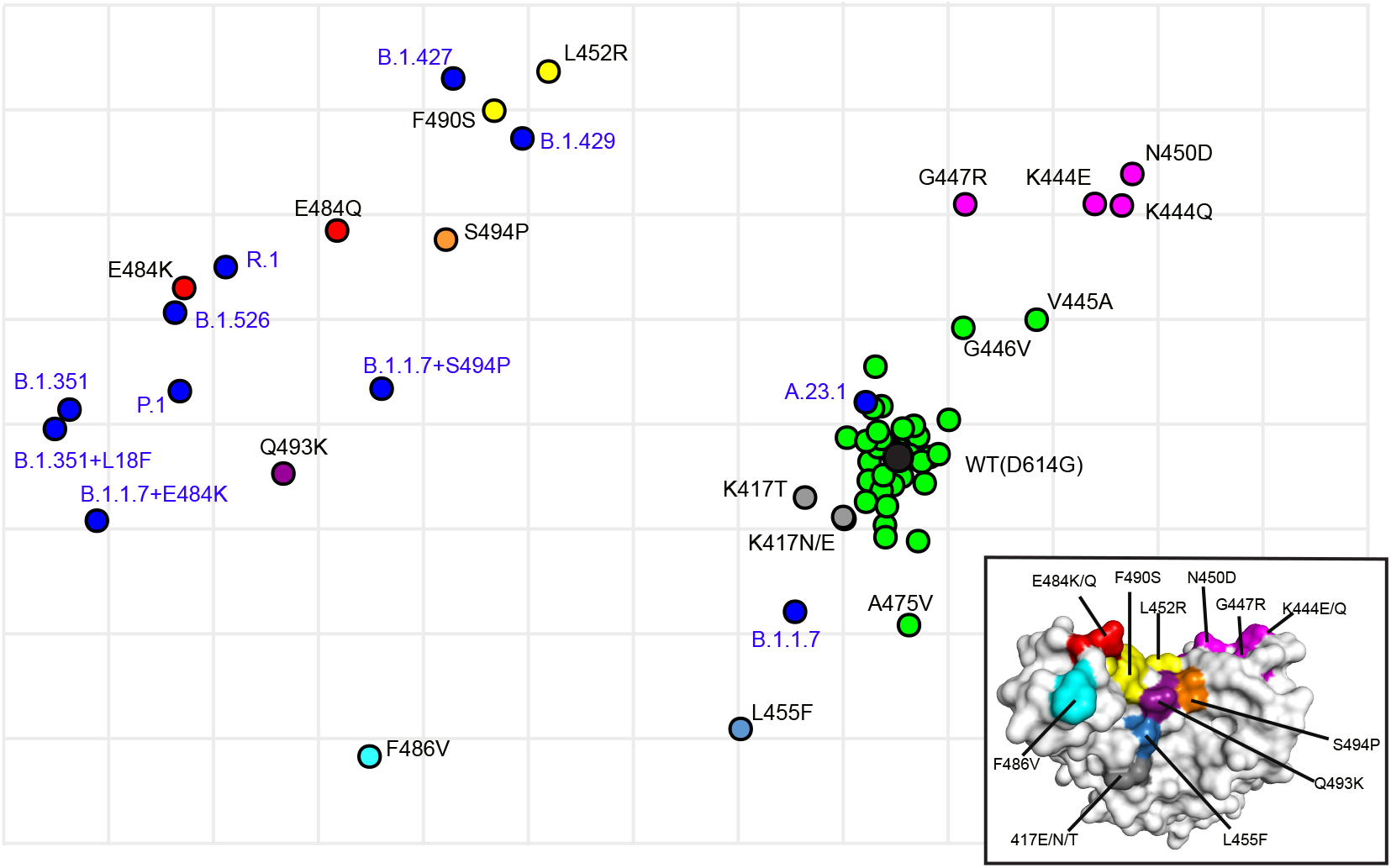
Antigenic cartography showing the relative antigenic distance of pseudoviruses with single substitutions in spike compared to spikes with the full set of substitutions found in variants. Antigenic maps were constructed using neutralization titers (dilution factors) of nAbs against all tested pseudoviruses. Blue dots identify pseudoviruses bearing Spikes representing variants of concern or interest. Black dot identifies the wild type (WT D614G) pseudovirus. Green dots identify pseudoviruses with single substitutions in Spike that are antigenically close to WT. Other colors identify pseudoviruses with single substitutions in Spike that are more antigenically distant from WT. Inset shows the color-coded locations of the single residue substitutions in the RBD.

## Discussion

This study compared the neutralization profiles of 25 nAbs, cnAbs, or pnAbs in late stages of clinical development or available under EUA against a panel of pseudoviruses bearing Spikes with single or multiple amino acid substitutions, including complete sets of mutations representing the B.1.1.7, B.1.351, P.1, B.1.427/B.1.429, B.1.526, R1, and A.23.1 variants. This large dataset confirms and extends results from other studies and highlights several issues that inform treatment strategies. We found that single amino acid substitutions in Spike were sufficient for conferring a loss of potency for 13 of 15 single nAbs, underscoring the importance of using combination nAbs with complementary resistance profiles. Six of the 15 single nAbs lost potency against E484K, which is present in multiple lineages. We also show that key substitutions in VOCs or VOIs can predict a loss of potency for susceptible nAbs, but other substitutions can modify the effects in unpredictable ways. These data highlight a role for epistatic interactions among residues within or outside the RBD that can affect antibody binding and potentially virus entry and evolution.

Among the 60 pseudoviruses with single or combinations of Spike substitutions all but three were functional (Fig S3), demonstrating the high degree of plasticity in Spike. Deep mutational scanning studies of the RBD previously showed that RBD binding to ACE2 can tolerate many substitutions (61). Spike plasticity facilitates virus adaption to new hosts and host tissues, as well as immune escape. Adaptive and immune selective pressures are problematic for nAbs selected from convalescent donors in early 2020, as the most potent antibodies were often against immune dominant epitopes involving the 484 position (15, 56, 62, 63). Clinical studies conducted as early as September 2020 indicated resistant variants were present at 6% in the untreated placebo group (64), which may indicate adaptive or immune pressure.

New variants are increasingly acquiring many Spike mutations. Here we identify key substitutions conferring resistance to clinical-stage nAbs and show different examples of resistance conferred by a single RBD substitution being modified by the context of other substitutions in the variant. For nAb.N, the resistance conferred by E484K was unexpectedly modified by the non-RBD substitution G769V or its combination with W152L (Fig. 4B and C, nAb. N bottom left panel and second panel, respectively), suggesting long-range epistatic interactions in Spike. We note that a G769E substitution was previously identified in an escape mutant study involving selection with RBD antibodies (65). In the case of nAb.J, resistance conferred by E484K alone was reversed in the context of P.1 and B.1.531, but not in the context of B.1.526 or R.1 (Fig 4B and C, nAb.J bottom panel and third panel, respectively). The B.1.526 and R1 variants only have the E484K substitution in the RBD, but B.1.351 and P.1 both have additional RBD substitutions at residues 417 and 501. NAb.J was also more potent against P.1 compared to B.1.351. Yet the potency of nAb.J was not affected by the single N501Y substitution, and the single K417N or K417T substitution in B.1.351 and P.1, respectively each enhanced nAb.J potency to a similar extent. These findings suggest epistatic interactions among residues 417, 484, and 501 in the conformationally dynamic RBD, consistent with their convergent evolution in independent virus lineages. The mechanistic details of how these residues may functionally interact needs further study. Epistatic networks are known to constrain the evolution of viral proteins. In the case of influenza, complex epistatic networks in the receptor binding site of influenza hemagglutinin have been reported to limit reversion of mutations (66), and epistatic interactions that maintain protein stability was suggested to constrain the evolution of the influenza nucleoprotein (67).

The continuing evolution of circulating SARS-CoV-2 variants, often containing substitutions known to confer resistance to several monoclonal antibodies, underscores the need for ongoing assessments of nAbs to ensure that they remain potent against new variants. nAbs in cocktails should have complementary resistance profiles and similar potency. However, the presence of MOCs, such as those leading to the E484 and L452 substitutions, in variants that are increasing in prevalence cautions against the use of nAbs vulnerable these substitutions. In a two nAb cocktail, a nAb resistant to E484K would effectively turn the cocktail into monotherapy against any E484K containing variant, rendering the cocktail less resilient against emerging substitutions. Additional nAbs with different resistance profiles in the RBD, as well as other regions in Spike, are therefore needed.

Finally, the safety and simplicity of pseudovirus neutralization assays offer many advantages for identifying MOCs and screening antibody therapeutics and vaccinee sera, but such assays have limitations. Pseudoviruses do not completely mimic authentic SARS-CoV-2 or virus entry conditions *in vivo*. In our nAb screens, we found that two antibodies that bind outside of the receptor binding motif could only partially neutralize (< 80%) pseudoviruses bearing WT Spike (data not shown). This inability to achieve complete neutralization by some nAbs has been observed by others using authentic SARS-CoV-2 (54, 68) and may be due to use of engineered target cells that permit efficient virus entry or heterogeneous glycosylation of Spike that affects an antibody epitope. Neutralization assays also cannot measure the potential protective activity afforded by nAb Fc effector functions. To address the limitations of any single assay, several consortia have formed to integrate findings from animal models and *in vitro* assays involving a variety of formats to inform the development of potent antibody therapeutics and other medical countermeasure that will remain potent against emerging variants.

## Methods

### Plasmids and cell lines

Codon-optimized full-length S genes of SARS-COV-2 variants (using D614G as the base plasmid) were cloned into pCDNA3.1(+) (GenScript, Piscataway, NJ). The HIV gag/pol (pCMVΔR8.2) and Luciferase reporter (pHR’CMV-Luc) plasmids were obtained from the Vaccine Research Center (National Institutes of Health, Bethesda, MD). 293T-ACE2/TMPRSS2 cells stably expressing ACE2 and transmembrane serine protease 2 (TMPRSS2) were established and maintained as previously described (51).

### Antibody therapeutics

Antibody therapeutics for COVID-19 were provided by pharmaceutical companies to support the US government COVID-19 therapeutics response team study to profile resistance of antibody therapeutic products.

### Choice of variants for study

An initial list of variants was generated by analyzing the GISAID dataset through October 2020. Selections were based on known nAb resistance from the literature, functional domains, and sample frequency. The most common NTD variants (global frequency > 0.2%) and all RBD variants observed in greater than 3 samples were included. Additional variants with multiple Spike mutations and their individual components were added as they were observed in GISAID. Priorities were calculated and re-evaluated weekly using four major variables: known nAb resistance (by number of nAbs affected), global frequency (% of global samples), increasing frequency trends over time (≥2-fold increase in a country with good sequence coverage in the most recent 4 weeks), and United States geographic spread (number of states with variant cases). The variants and accession numbers for the spike sequences used in this study are listed in Figure S1.

### SARS-CoV-2 pseudovirus production and neutralization assay

Lentiviral pseudoviruses bearing Spikes of variants were generated and used for neutralization studies in stable 293T-ACE2/TMPRSS2 cells (BEI # NR-55351) as previously described (51). Pseudoviruses with titers of approximately 10^6^ relative luminescence units (RLU)/ml were incubated with four-fold serially-diluted antibodies for two hr at 37°C prior to inoculation onto target cells in 96-well plates and scoring for luminescence activity 48 hr later. Titers were calculated using a nonlinear regression curve fit (GraphPad Prism software Inc., La Jolla, CA). The nAb concentration or inverse of the dilutions causing a 50% reduction of RLU compared to control (IC_50_) was reported as the neutralizing antibody titer. Antibodies not reaching 80% neutralization were considered resistant (R), and IC_50_ values were reported as greater than the highest concentration tested. The mean titer from at least two independent experiments each with intra-assay duplicates was reported as the final titer. WT was run as a control for each assay.

### Antigenic Cartography

We created a geometric interpretation of neutralization titers against the tested SARS-CoV-2 pseudoviruses using ACMAS antigenic cartography software (https://acmacs-web.antigenic-cartography.org/). The map is presented on a grid in which each square indicates one antigenic unit, corresponding to a two-fold dilution of the antibody in the neutralization assay. Antigenic distance is measured in any direction on the map.

### Computational analysis

The amino acid locations on the RBD that affected nAb potency were modeled with PyMOL molecular graphics system (version 1.7.4; Schrödinger, LLC) using the Protein Data Bank (PDB) entries 6ZOZ for the closed conformation and 7A98 for the open conformation.

### Statistics analysis

Statistical significance was determined by the two-tailed, unpaired t test using GraphPad Prism 8 software (San Diego, CA).

## Abbreviations

(SARS-CoV-2): severe acute respiratory syndrome coronavirus 2
(COVID-19): coronavirus disease 2019
(neutralizing antibodies): nAbs
(Spike): coronavirus spike glycoprotein
(RBD): receptor binding domain
(NTD): N-terminal domain
(EUA): emergency use authorization
(ACE2): angiotension converting enzyme 2
(TMPRSS2): transmembrane protease serine 2 precursor
(S1): spike surface subunit
(S2): spike transmembrane subunit
(VOC): variant of concern
(VOI): variant of interest
(MOC): mutation of concern

## Acknowledgements

We thank Drs. Maryna Eichelberger and Gabriel Parra (US Food and Drug Administration, Centers for Biologics Evaluation and Research) for critical review of the manuscript.

---

**Figure S1.**
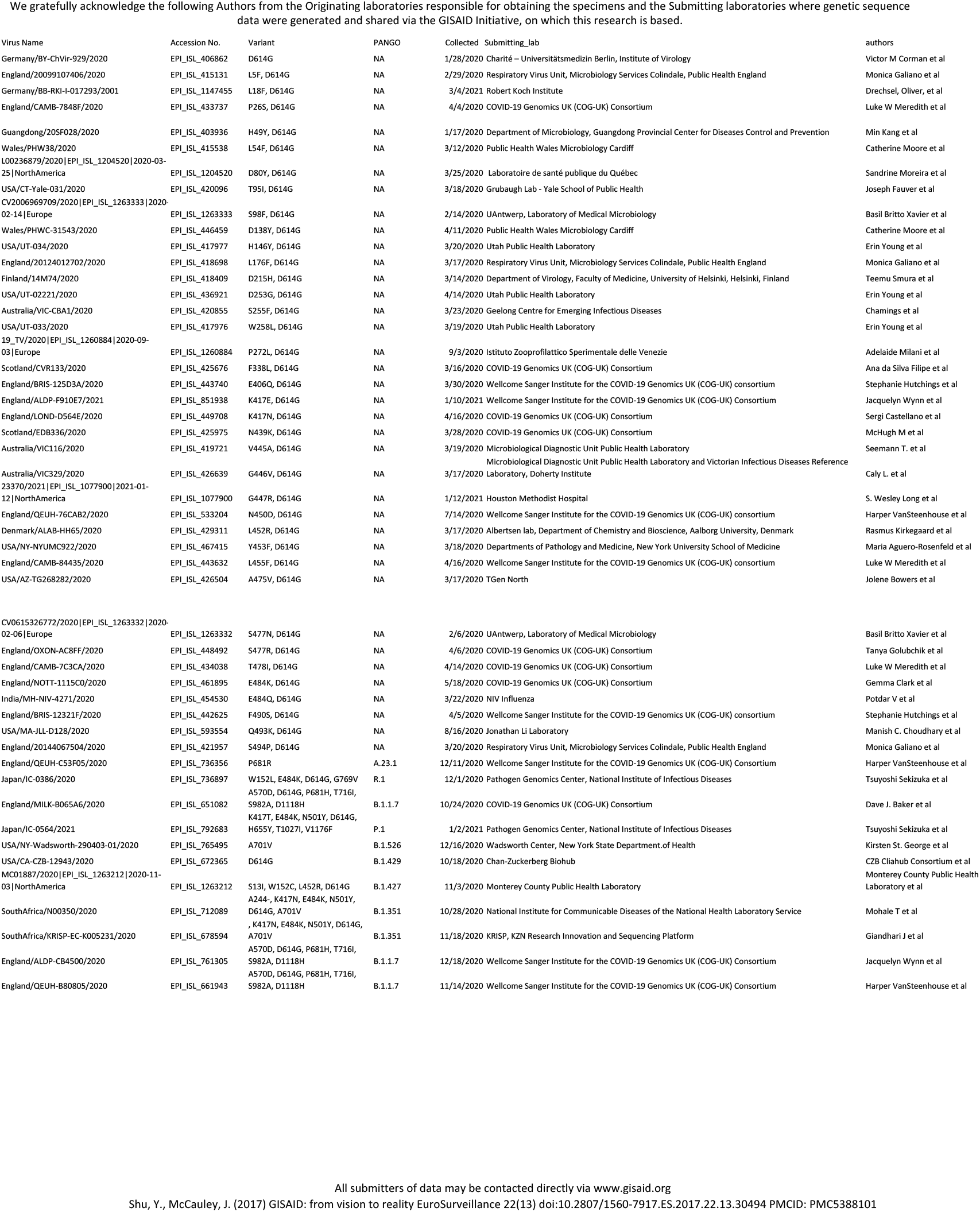
List of the variants from GISAID that were used in this study.

**Figure S2.**
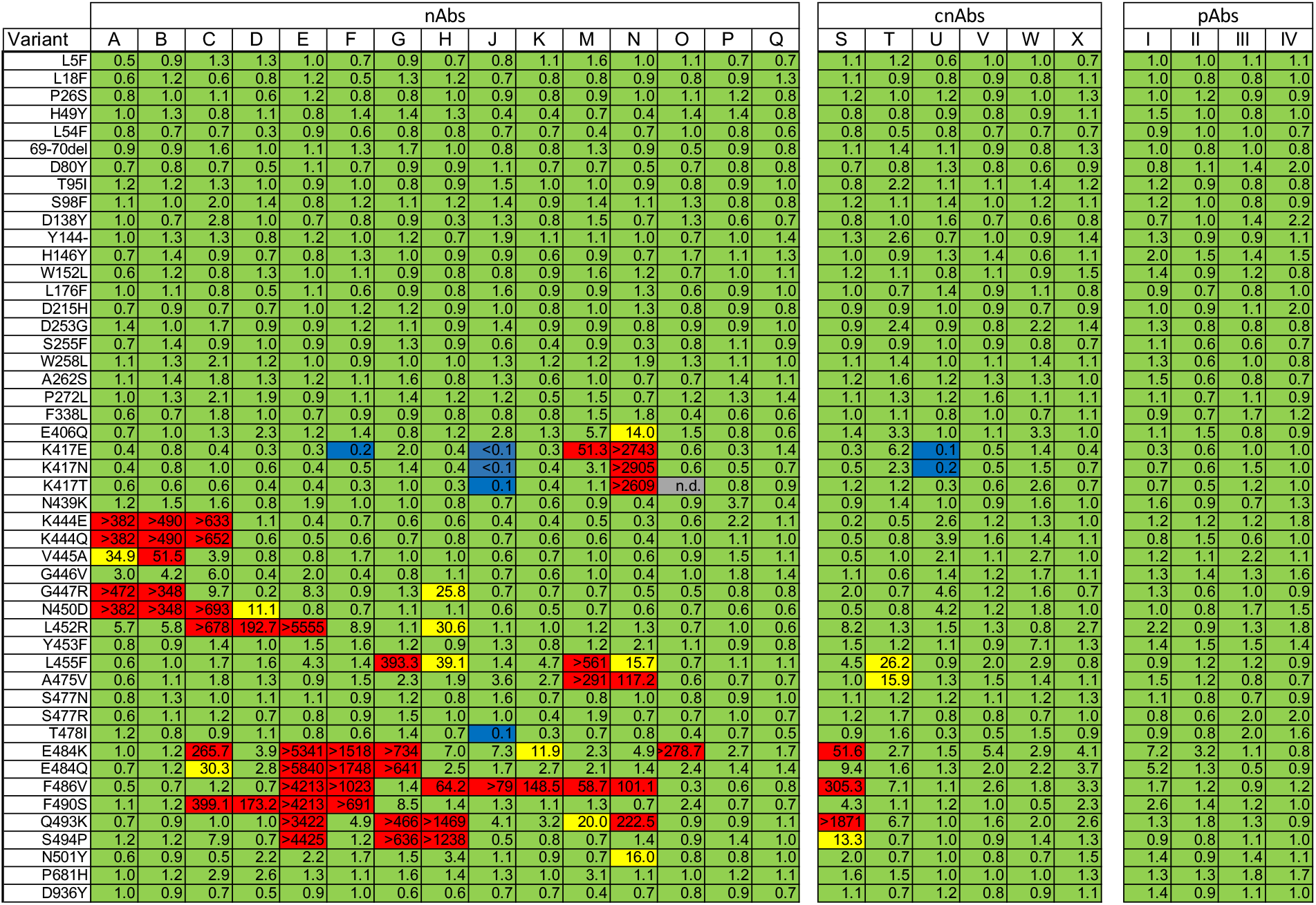
Heat map showing nAb potency as the ratio between the IC_50_ values of pseudoviruses with single substitutions or deletions against IC_50_ of WT pseudoviruses for each antibody. Ratios are color coded as follows: <10, green = no major impact; 10-50 yellow = moderate loss of potency and >50 red = highly resistant. <0.2, blue = increased potency. Combinations not determined are shown in gray (n.d.) Data shown represent at least two independent experiments each with an intra-assay duplicate, except where indicated by *. * indicates only tested once.

**Figure S3.**
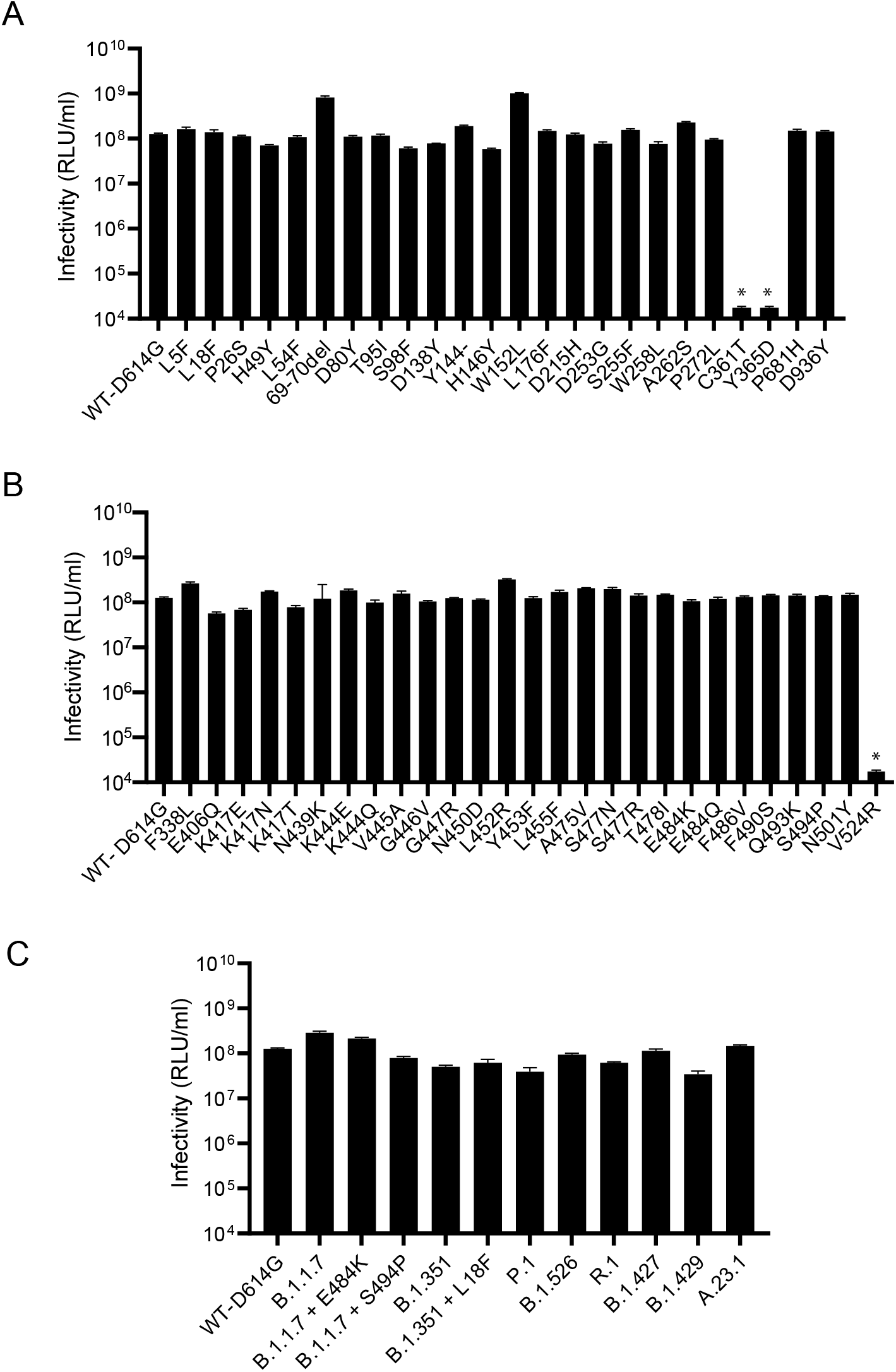
Infectivity of the pseudoviruses bearing Spikes with the single and multiple amino acid substitutions. (A) Infectivity of pseudoviruses with deletions (del) or single substitutions in the NTD and S2 regions of Spike. (B) Infectivity of pseudoviruses with single substitutions in the RBD. (C) Infectivity of pseudoviruses with multiple substitutions representing VOCs or VOIs. Volume-normalized transfection supernatants were used to inoculate stable 293T-ACE2/TMPRSS2 cells. Results shown are averages of two independent experiments. Infectivity expressed as relative luminescence units per ml (RLU/ml) with standard deviation. Data shown represent at least two independent experiments each with an intra-assay duplicate. * indicates pseudoviruses with low infectivity that could not be used for neutralization studies.

**Table S1.**
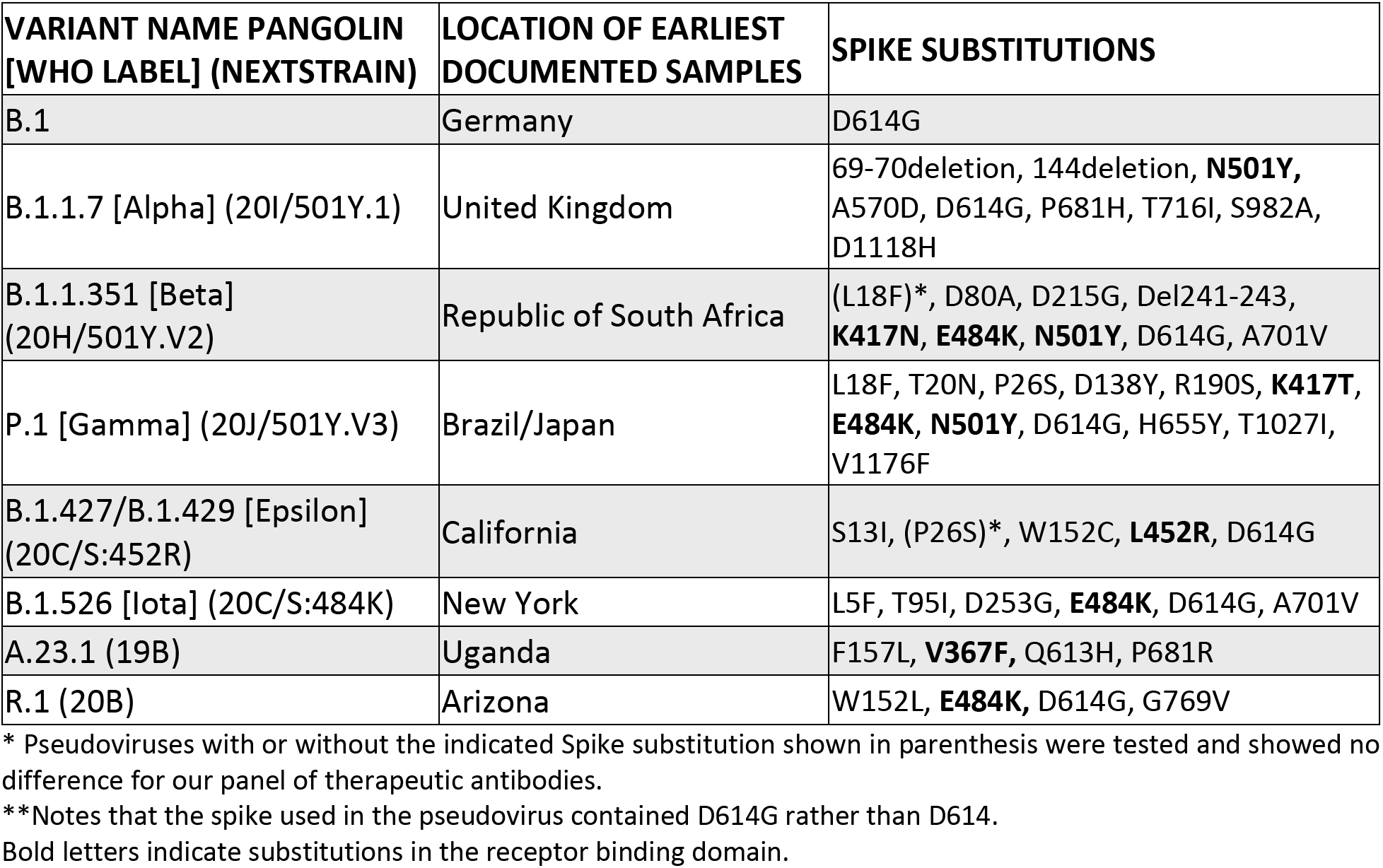
SARS-CoV-2 Variants.

