## Supplementary material for "Key substitutions in the spike protein of SARS-CoV-2 variants can predict resistance to monoclonal antibodies, but other substitutions can modify the effects"

We gratefully acknowledge the following Authors from the Originating laboratories responsible for obtaining the specimens and the Submitting laboratories where genetic sequence data were generated and shared via the GISAID Initiative, on which this research is based.

| Virus Name | Accession No. | Variant | PANGO | Collected | Submitting_lab | authors |
|---|---|---|---|---|---|---|
| Germany/BY-ChVir-929/2020 | EPI_ISL_406862 | D614G | NA | 1/28/2020 | Charité – Universitätsmedizin Berlin, Institute of Virology | Victor M Corman et al |
| England/20099107406/2020 | EPI_ISL_415131 | L5F, D614G | NA | 2/29/2020 | Respiratory Virus Unit, Microbiology Services Colindale, Public Health England | Monica Galiano et al |
| Germany/BB-RKI-I-017293/2001 | EPI_ISL_1147455 | L18F, D614G | NA | 3/4/2021 | Robert Koch Institute | Drechsel, Oliver, et al |
| England/CAMB-7848F/2020 | EPI_ISL_433737 | P26S, D614G | NA | 4/4/2020 | COVID-19 Genomics UK (COG-UK) Consortium | Luke W Meredith et al |
| Guangdong/20SF028/2020 | EPI_ISL_403936 | H49Y, D614G | NA | 1/17/2020 | Department of Microbiology, Guangdong Provincial Center for Diseases Control and Prevention | Min Kang et al |
| Wales/PHW38/2020 | EPI_ISL_415538 | L54F, D614G | NA | 3/12/2020 | Public Health Wales Microbiology Cardiff | Catherine Moore et al |
| L00236879/2020\|EPI_ISL_1204520\|2020-03-25\|NorthAmerica | EPI_ISL_1204520 | D80Y, D614G | NA | 3/25/2020 | Laboratoire de santé publique du Québec | Sandrine Moreira et al |
| USA/CT-Yale-031/2020 | EPI_ISL_420096 | T95I, D614G | NA | 3/18/2020 | Grubaugh Lab - Yale School of Public Health | Joseph Fauver et al |
| CV2006969709/2020\|EPI_ISL_1263333\|2020-02-14\|Europe | EPI_ISL_1263333 | S98F, D614G | NA | 2/14/2020 | UAntwerp, Laboratory of Medical Microbiology | Basil Britto Xavier et al |
| Wales/PHWC-31543/2020 | EPI_ISL_446459 | D138Y, D614G | NA | 4/11/2020 | Public Health Wales Microbiology Cardiff | Catherine Moore et al |
| USA/UT-034/2020 | EPI_ISL_417977 | H146Y, D614G | NA | 3/20/2020 | Utah Public Health Laboratory | Erin Young et al |
| England/20124012702/2020 | EPI_ISL_418698 | L176F, D614G | NA | 3/17/2020 | Respiratory Virus Unit, Microbiology Services Colindale, Public Health England | Monica Galiano et al |
| Finland/14M74/2020 | EPI_ISL_418409 | D215H, D614G | NA | 3/14/2020 | Department of Virology, Faculty of Medicine, University of Helsinki, Helsinki, Finland | Teemu Smura et al |
| USA/UT-02221/2020 | EPI_ISL_436921 | D253G, D614G | NA | 4/14/2020 | Utah Public Health Laboratory | Erin Young et al |
| Australia/VIC-CBA1/2020 | EPI_ISL_420855 | S255F, D614G | NA | 3/23/2020 | Geelong Centre for Emerging Infectious Diseases | Chamings et al |
| USA/UT-033/2020 | EPI_ISL_417976 | W258L, D614G | NA | 3/19/2020 | Utah Public Health Laboratory | Erin Young et al |
| 19_TV/2020\|EPI_ISL_1260884\|2020-09-03\|Europe | EPI_ISL_1260884 | P272L, D614G | NA | 9/3/2020 | Istituto Zooprofilattico Sperimentale delle Venezie | Adelaide Milani et al |
| Scotland/CVR133/2020 | EPI_ISL_425676 | F338L, D614G | NA | 3/16/2020 | COVID-19 Genomics UK (COG-UK) Consortium | Ana da Silva Filipe et al |
| England/BRIS-125D3A/2020 | EPI_ISL_443740 | E406Q, D614G | NA | 3/30/2020 | Wellcome Sanger Institute for the COVID-19 Genomics UK (COG-UK) consortium | Stephanie Hutchings et al |
| England/ALDP-F910E7/2021 | EPI_ISL_851938 | K417E, D614G | NA | 1/10/2021 | Wellcome Sanger Institute for the COVID-19 Genomics UK (COG-UK) Consortium | Jacquelyn Wynn et al |
| England/LOND-D564E/2020 | EPI_ISL_449708 | K417N, D614G | NA | 4/16/2020 | COVID-19 Genomics UK (COG-UK) Consortium | Sergi Castellano et al |
| Scotland/EDB336/2020 | EPI_ISL_425975 | N439K, D614G | NA | 3/28/2020 | COVID-19 Genomics UK (COG-UK) Consortium | McHugh M et al |
| Australia/VIC116/2020 | EPI_ISL_419721 | V445A, D614G | NA | 3/19/2020 | Microbiological Diagnostic Unit Public Health Laboratory | Seemann T. et al |
| Australia/VIC329/2020 | EPI_ISL_426639 | G446V, D614G | NA | 3/17/2020 | Microbiological Diagnostic Unit Public Health Laboratory and Victorian Infectious Diseases Reference Laboratory, Doherty Institute | Caly L. et al |
| 23370/2021\|EPI_ISL_1077900\|2021-01-12\|NorthAmerica | EPI_ISL_1077900 | G447R, D614G | NA | 1/12/2021 | Houston Methodist Hospital | S. Wesley Long et al |
| England/QEUH-76CAB2/2020 | EPI_ISL_533204 | N450D, D614G | NA | 7/14/2020 | Wellcome Sanger Institute for the COVID-19 Genomics UK (COG-UK) consortium | Harper VanSteenhouse et al |
| Denmark/ALAB-HH65/2020 | EPI_ISL_429311 | L452R, D614G | NA | 3/17/2020 | Albertsen lab, Department of Chemistry and Bioscience, Aalborg University, Denmark | Rasmus Kirkegaard et al |
| USA/NY-NYUMC922/2020 | EPI_ISL_467415 | Y453F, D614G | NA | 3/18/2020 | Departments of Pathology and Medicine, New York University School of Medicine | Maria Aguero-Rosenfeld et al |
| England/CAMB-84435/2020 | EPI_ISL_443632 | L455F, D614G | NA | 4/16/2020 | Wellcome Sanger Institute for the COVID-19 Genomics UK (COG-UK) consortium | Luke W Meredith et al |
| USA/AZ-TG268282/2020 | EPI_ISL_426504 | A475V, D614G | NA | 3/17/2020 | TGen North | Jolene Bowers et al |
| CV0615326772/2020\|EPI_ISL_1263332\|2020-02-06\|Europe | EPI_ISL_1263332 | S477N, D614G | NA | 2/6/2020 | UAntwerp, Laboratory of Medical Microbiology | Basil Britto Xavier et al |
| England/OXON-AC8FF/2020 | EPI_ISL_448492 | S477R, D614G | NA | 4/6/2020 | COVID-19 Genomics UK (COG-UK) Consortium | Tanya Golubchik et al |
| England/CAMB-7C3CA/2020 | EPI_ISL_434038 | T478I, D614G | NA | 4/14/2020 | COVID-19 Genomics UK (COG-UK) Consortium | Luke W Meredith et al |
| England/NOTT-1115C0/2020 | EPI_ISL_461895 | E484K, D614G | NA | 5/18/2020 | COVID-19 Genomics UK (COG-UK) Consortium | Gemma Clark et al |
| India/MH-NIV-4271/2020 | EPI_ISL_454530 | E484Q, D614G | NA | 3/22/2020 | NIV Influenza | Potdar V et al |
| England/BRIS-12321F/2020 | EPI_ISL_442625 | F490S, D614G | NA | 4/5/2020 | Wellcome Sanger Institute for the COVID-19 Genomics UK (COG-UK) consortium | Stephanie Hutchings et al |
| USA/MA-JLL-D128/2020 | EPI_ISL_593554 | Q493K, D614G | NA | 8/16/2020 | Jonathan Li Laboratory | Manish C. Choudhary et al |
| England/20144067504/2020 | EPI_ISL_421957 | S494P, D614G | NA | 3/20/2020 | Respiratory Virus Unit, Microbiology Services Colindale, Public Health England | Monica Galiano et al |
| England/QEUH-C53F05/2020 | EPI_ISL_736356 | P681R | A.23.1 | 12/11/2020 | Wellcome Sanger Institute for the COVID-19 Genomics UK (COG-UK) Consortium | Harper VanSteenhouse et al |
| Japan/IC-0386/2020 | EPI_ISL_736897 | W152L, E484K, D614G, G769V | R.1 | 12/1/2020 | Pathogen Genomics Center, National Institute of Infectious Diseases | Tsuyoshi Sekizuka et al |
| England/MILK-B065A6/2020 | EPI_ISL_651082 | A570D, D614G, P681H, T716I, S982A, D1118H | B.1.1.7 | 10/24/2020 | COVID-19 Genomics UK (COG-UK) Consortium | Dave J. Baker et al |
| Japan/IC-0564/2021 | EPI_ISL_792683 | K417T, E484K, N501Y, D614G, H655Y, T1027I, V1176F | P.1 | 1/2/2021 | Pathogen Genomics Center, National Institute of Infectious Diseases | Tsuyoshi Sekizuka et al |
| USA/NY-Wadsworth-290403-01/2020 | EPI_ISL_765495 | A701V | B.1.526 | 12/16/2020 | Wadsworth Center, New York State Department.of Health | Kirsten St. George et al |
| USA/CA-CZB-12943/2020 | EPI_ISL_672365 | D614G | B.1.429 | 10/18/2020 | Chan-Zuckerberg Biohub | CZB Cliahub Consortium et al |
| MC01887/2020\|EPI_ISL_1263212\|2020-11-03\|NorthAmerica | EPI_ISL_1263212 | S13I, W152C, L452R, D614G | B.1.427 | 11/3/2020 | Monterey County Public Health Laboratory | Monterey County Public Health Laboratory et al |
| SouthAfrica/N00350/2020 | EPI_ISL_712089 | A244-, K417N, E484K, N501Y, D614G, A701V | B.1.351 | 10/28/2020 | National Institute for Communicable Diseases of the National Health Laboratory Service | Mohale T et al |
| SouthAfrica/KRISP-EC-K005231/2020 | EPI_ISL_678594 | , K417N, E484K, N501Y, D614G, A701V | B.1.351 | 11/18/2020 | KRISP, KZN Research Innovation and Sequencing Platform | Giandhari J et al |
| England/ALDP-CB4500/2020 | EPI_ISL_761305 | A570D, D614G, P681H, T716I, S982A, D1118H | B.1.1.7 | 12/18/2020 | Wellcome Sanger Institute for the COVID-19 Genomics UK (COG-UK) Consortium | Jacquelyn Wynn et al |
| England/QEUH-B80805/2020 | EPI_ISL_661943 | A570D, D614G, P681H, T716I, S982A, D1118H | B.1.1.7 | 11/14/2020 | Wellcome Sanger Institute for the COVID-19 Genomics UK (COG-UK) Consortium | Harper VanSteenhouse et al |

All submitters of data may be contacted directly via www.gisaid.org

Shu, Y., McCauley, J. (2017) GISAID: from vision to reality EuroSurveillance 22(13) doi:10.2807/1560-7917.ES.2017.22.13.30494 PMCID: PMC5388101

**Figure S1. List of the variants from GISAID that were used in this study.**

| | nAbs | | | | | | | | | | | | | | | cnAbs | | | | | | pAbs | | | |
|---|---|---|---|---|---|---|---|---|---|---|---|---|---|---|---|---|---|---|---|---|---|---|---|---|---|
| Variant | A | B | C | D | E | F | G | H | J | K | M | N | O | P | Q | S | T | U | V | W | X | I | II | III | IV |
| L5F | 0.5 | 0.9 | 1.3 | 1.3 | 1.0 | 0.7 | 0.9 | 0.7 | 0.8 | 1.1 | 1.6 | 1.0 | 1.1 | 0.7 | 0.7 | 1.1 | 1.2 | 0.6 | 1.0 | 1.0 | 0.7 | 1.0 | 1.0 | 1.1 | 1.1 |
| L18F | 0.6 | 1.2 | 0.6 | 0.8 | 1.2 | 0.5 | 1.3 | 1.2 | 0.7 | 0.8 | 0.8 | 0.9 | 0.8 | 0.9 | 1.3 | 1.1 | 0.9 | 0.8 | 0.9 | 0.8 | 1.1 | 1.0 | 0.8 | 0.8 | 1.0 |
| P26S | 0.8 | 1.0 | 1.1 | 0.6 | 1.2 | 0.8 | 0.8 | 1.0 | 0.9 | 0.8 | 0.9 | 1.0 | 1.1 | 1.2 | 0.8 | 1.2 | 1.0 | 1.2 | 0.9 | 1.0 | 1.3 | 1.0 | 1.2 | 0.9 | 0.9 |
| H49Y | 1.0 | 1.3 | 0.8 | 1.1 | 0.8 | 1.4 | 1.4 | 1.3 | 0.4 | 0.4 | 0.7 | 0.4 | 1.4 | 1.4 | 0.8 | 0.8 | 0.8 | 0.9 | 0.8 | 0.9 | 1.1 | 1.5 | 1.0 | 0.8 | 1.0 |
| L54F | 0.8 | 0.7 | 0.7 | 0.3 | 0.9 | 0.6 | 0.8 | 0.8 | 0.7 | 0.7 | 0.4 | 0.7 | 1.0 | 0.8 | 0.6 | 0.8 | 0.5 | 0.8 | 0.7 | 0.7 | 0.7 | 0.9 | 1.0 | 1.0 | 0.7 |
| 69-70del | 0.9 | 0.9 | 1.6 | 1.0 | 1.1 | 1.3 | 1.7 | 1.0 | 0.8 | 0.8 | 1.3 | 0.9 | 0.5 | 0.9 | 0.8 | 1.1 | 1.4 | 1.1 | 0.9 | 0.8 | 1.3 | 1.0 | 0.8 | 1.0 | 0.8 |
| D80Y | 0.7 | 0.8 | 0.7 | 0.5 | 1.1 | 0.7 | 0.9 | 0.9 | 1.1 | 0.7 | 0.7 | 0.5 | 0.7 | 0.8 | 0.8 | 0.7 | 0.8 | 1.3 | 0.8 | 0.6 | 0.9 | 0.8 | 1.1 | 1.4 | 2.0 |
| T95I | 1.2 | 1.2 | 1.3 | 1.0 | 0.9 | 1.0 | 0.8 | 0.9 | 1.5 | 1.0 | 1.0 | 0.9 | 0.8 | 0.9 | 1.0 | 0.8 | 2.2 | 1.1 | 1.1 | 1.4 | 1.2 | 1.2 | 0.9 | 0.8 | 0.8 |
| S98F | 1.1 | 1.0 | 2.0 | 1.4 | 0.8 | 1.2 | 1.1 | 1.2 | 1.4 | 0.9 | 1.4 | 1.1 | 1.3 | 0.8 | 0.8 | 1.2 | 1.1 | 1.4 | 1.0 | 1.2 | 1.1 | 1.2 | 1.0 | 0.8 | 0.9 |
| D138Y | 1.0 | 0.7 | 2.8 | 1.0 | 0.7 | 0.8 | 0.9 | 0.3 | 1.3 | 0.8 | 1.5 | 0.7 | 1.3 | 0.6 | 0.7 | 0.8 | 1.0 | 1.6 | 0.7 | 0.6 | 0.8 | 0.7 | 1.0 | 1.4 | 2.2 |
| Y144- | 1.0 | 1.3 | 1.3 | 0.8 | 1.2 | 1.0 | 1.2 | 0.7 | 1.9 | 1.1 | 1.1 | 1.0 | 0.7 | 1.0 | 1.4 | 1.3 | 2.6 | 0.7 | 1.0 | 0.9 | 1.4 | 1.3 | 0.9 | 0.9 | 1.1 |
| H146Y | 0.7 | 1.4 | 0.9 | 0.7 | 0.8 | 1.3 | 1.0 | 0.9 | 0.9 | 0.6 | 0.9 | 0.7 | 1.7 | 1.1 | 1.3 | 1.0 | 0.9 | 1.3 | 1.4 | 0.6 | 1.1 | 2.0 | 1.5 | 1.4 | 1.5 |
| W152L | 0.6 | 1.2 | 0.8 | 1.3 | 1.0 | 1.1 | 0.9 | 0.8 | 0.8 | 0.9 | 1.6 | 1.2 | 0.7 | 1.0 | 1.1 | 1.2 | 1.1 | 0.8 | 1.1 | 0.9 | 1.5 | 1.4 | 0.9 | 1.2 | 0.8 |
| L176F | 1.0 | 1.1 | 0.8 | 0.5 | 1.1 | 0.6 | 0.9 | 0.8 | 1.6 | 0.9 | 0.9 | 1.3 | 0.6 | 0.9 | 1.0 | 1.0 | 0.7 | 1.4 | 0.9 | 1.1 | 0.8 | 0.9 | 0.7 | 0.8 | 1.0 |
| D215H | 0.7 | 0.9 | 0.7 | 0.7 | 1.0 | 1.2 | 1.2 | 0.9 | 1.0 | 0.8 | 1.0 | 1.3 | 0.8 | 0.9 | 0.8 | 0.9 | 0.9 | 1.0 | 0.9 | 0.7 | 0.9 | 1.0 | 0.9 | 1.1 | 2.0 |
| D253G | 1.4 | 1.0 | 1.7 | 0.9 | 0.9 | 1.2 | 1.1 | 0.9 | 1.4 | 0.9 | 0.9 | 0.8 | 0.9 | 0.9 | 1.0 | 0.9 | 2.4 | 0.9 | 0.8 | 2.2 | 1.4 | 1.3 | 0.8 | 0.8 | 0.8 |
| S255F | 0.7 | 1.4 | 0.9 | 1.0 | 0.9 | 0.9 | 1.3 | 0.9 | 0.6 | 0.4 | 0.9 | 0.3 | 0.8 | 1.1 | 0.9 | 0.9 | 0.9 | 1.0 | 0.9 | 0.8 | 0.7 | 1.1 | 0.6 | 0.6 | 0.7 |
| W258L | 1.1 | 1.3 | 2.1 | 1.2 | 1.0 | 0.9 | 1.0 | 1.0 | 1.3 | 1.2 | 1.2 | 1.9 | 1.3 | 1.1 | 1.0 | 1.1 | 1.4 | 1.0 | 1.1 | 1.4 | 1.1 | 1.3 | 0.6 | 1.0 | 0.8 |
| A262S | 1.1 | 1.4 | 1.8 | 1.3 | 1.2 | 1.1 | 1.6 | 0.8 | 1.3 | 0.6 | 1.0 | 0.7 | 0.7 | 1.4 | 1.1 | 1.2 | 1.6 | 1.2 | 1.3 | 1.3 | 1.0 | 1.5 | 0.6 | 0.8 | 0.7 |
| P272L | 1.0 | 1.3 | 2.1 | 1.9 | 0.9 | 1.1 | 1.4 | 1.2 | 1.2 | 0.5 | 1.5 | 0.7 | 1.2 | 1.3 | 1.4 | 1.1 | 1.3 | 1.2 | 1.6 | 1.1 | 1.1 | 1.1 | 0.7 | 1.1 | 0.9 |
| F338L | 0.6 | 0.7 | 1.8 | 1.0 | 0.7 | 0.9 | 0.9 | 0.8 | 0.8 | 0.8 | 1.5 | 1.8 | 0.4 | 0.6 | 0.6 | 1.0 | 1.1 | 0.8 | 1.0 | 0.7 | 1.1 | 0.9 | 0.7 | 1.7 | 1.2 |
| E406Q | 0.7 | 1.0 | 1.3 | 2.3 | 1.2 | 1.4 | 0.8 | 1.2 | 2.8 | 1.3 | 5.7 | 14.0 | 1.5 | 0.8 | 0.6 | 1.4 | 3.3 | 1.0 | 1.1 | 3.3 | 1.0 | 1.1 | 1.5 | 0.8 | 0.9 |
| K417E | 0.4 | 0.8 | 0.4 | 0.3 | 0.3 | 0.2 | 2.0 | 0.4 | <0.1 | 0.3 | 51.3 | >2743 | 0.6 | 0.3 | 1.4 | 0.3 | 6.2 | 0.1 | 0.5 | 1.4 | 0.4 | 0.3 | 0.6 | 1.0 | 1.0 |
| K417N | 0.4 | 0.8 | 1.0 | 0.6 | 0.4 | 0.5 | 1.4 | 0.4 | <0.1 | 0.4 | 3.1 | >2905 | 0.6 | 0.5 | 0.7 | 0.5 | 2.3 | 0.2 | 0.5 | 1.5 | 0.7 | 0.7 | 0.6 | 1.5 | 1.0 |
| K417T | 0.6 | 0.6 | 0.6 | 0.4 | 0.4 | 0.3 | 1.0 | 0.3 | 0.1 | 0.4 | 1.1 | >2609 | n.d. | 0.8 | 0.9 | 1.2 | 1.2 | 0.3 | 0.6 | 2.6 | 0.7 | 0.7 | 0.5 | 1.2 | 1.0 |
| N439K | 1.2 | 1.5 | 1.6 | 0.8 | 1.9 | 1.0 | 1.0 | 0.8 | 0.7 | 0.6 | 0.9 | 0.4 | 0.9 | 3.7 | 0.4 | 0.9 | 1.4 | 1.0 | 0.9 | 1.6 | 1.0 | 1.6 | 0.9 | 0.7 | 1.3 |
| K444E | >382 | >490 | >633 | 1.1 | 0.4 | 0.7 | 0.6 | 0.6 | 0.4 | 0.4 | 0.5 | 0.3 | 0.6 | 2.2 | 1.1 | 0.2 | 0.5 | 2.6 | 1.2 | 1.3 | 1.0 | 1.2 | 1.2 | 1.2 | 1.8 |
| K444Q | >382 | >490 | >652 | 0.6 | 0.5 | 0.6 | 0.7 | 0.8 | 0.7 | 0.6 | 0.6 | 0.4 | 1.0 | 1.1 | 1.0 | 0.5 | 0.8 | 3.9 | 1.6 | 1.4 | 1.1 | 0.8 | 1.5 | 0.6 | 1.0 |
| V445A | 34.9 | 51.5 | 3.9 | 0.8 | 0.8 | 1.7 | 1.0 | 1.0 | 0.6 | 0.7 | 1.0 | 0.6 | 0.9 | 1.5 | 1.1 | 0.5 | 1.0 | 2.1 | 1.1 | 2.7 | 1.0 | 1.2 | 1.1 | 2.2 | 1.1 |
| G446V | 3.0 | 4.2 | 6.0 | 0.4 | 2.0 | 0.4 | 0.8 | 1.1 | 0.7 | 0.6 | 1.0 | 0.4 | 1.0 | 1.8 | 1.4 | 1.1 | 0.6 | 1.4 | 1.2 | 1.7 | 1.1 | 1.3 | 1.4 | 1.3 | 1.6 |
| G447R | >472 | >348 | 9.7 | 0.2 | 8.3 | 0.9 | 1.3 | 25.8 | 0.7 | 0.7 | 0.7 | 0.5 | 0.5 | 0.8 | 0.8 | 2.0 | 0.7 | 4.6 | 1.2 | 1.6 | 0.7 | 1.3 | 0.6 | 1.0 | 0.9 |
| N450D | >382 | >348 | >693 | 11.1 | 0.8 | 0.7 | 1.1 | 1.1 | 0.6 | 0.5 | 0.7 | 0.6 | 0.7 | 0.6 | 0.6 | 0.5 | 0.8 | 4.2 | 1.2 | 1.8 | 1.0 | 1.0 | 0.8 | 1.7 | 1.5 |
| L452R | 5.7 | 5.8 | >678 | 192.7 | >5555 | 8.9 | 1.1 | 30.6 | 1.1 | 1.0 | 1.2 | 1.3 | 0.7 | 1.0 | 0.6 | 8.2 | 1.3 | 1.5 | 1.3 | 0.8 | 2.7 | 2.2 | 0.9 | 1.3 | 1.8 |
| Y453F | 0.8 | 0.9 | 1.4 | 1.0 | 1.5 | 1.6 | 1.2 | 0.9 | 1.3 | 0.8 | 1.2 | 2.1 | 1.1 | 0.9 | 0.8 | 1.5 | 1.2 | 1.1 | 0.9 | 7.1 | 1.3 | 1.4 | 1.5 | 1.5 | 1.4 |
| L455F | 0.6 | 1.0 | 1.7 | 1.6 | 4.3 | 1.4 | 393.3 | 39.1 | 1.4 | 4.7 | >561 | 15.7 | 0.7 | 1.1 | 1.1 | 4.5 | 26.2 | 0.9 | 2.0 | 2.9 | 0.8 | 0.9 | 1.2 | 1.2 | 0.9 |
| A475V | 0.6 | 1.1 | 1.8 | 1.3 | 0.9 | 1.5 | 2.3 | 1.9 | 3.6 | 2.7 | >291 | 117.2 | 0.6 | 0.7 | 0.7 | 1.0 | 15.9 | 1.3 | 1.5 | 1.4 | 1.1 | 1.5 | 1.2 | 0.8 | 0.7 |
| S477N | 0.8 | 1.3 | 1.0 | 1.1 | 1.1 | 0.9 | 1.2 | 0.8 | 1.6 | 0.7 | 0.8 | 1.0 | 0.8 | 0.9 | 1.0 | 1.1 | 1.2 | 1.2 | 1.1 | 1.2 | 1.3 | 1.1 | 0.8 | 0.7 | 0.9 |
| S477R | 0.6 | 1.1 | 1.2 | 0.7 | 0.8 | 0.9 | 1.5 | 1.0 | 1.0 | 0.4 | 1.9 | 0.7 | 0.7 | 1.0 | 0.7 | 1.2 | 1.7 | 0.8 | 0.8 | 0.7 | 1.0 | 0.8 | 0.6 | 2.0 | 2.0 |
| T478I | 1.2 | 0.8 | 0.9 | 1.1 | 0.8 | 0.6 | 1.4 | 0.7 | 0.1 | 0.3 | 0.7 | 0.8 | 0.4 | 0.7 | 0.5 | 0.9 | 1.6 | 0.3 | 0.5 | 1.5 | 0.9 | 0.9 | 0.8 | 2.0 | 1.6 |
| E484K | 1.0 | 1.2 | 265.7 | 3.9 | >5341 | >1518 | >734 | 7.0 | 7.3 | 11.9 | 2.3 | 4.9 | >278.7 | 2.7 | 1.7 | 51.6 | 2.7 | 1.5 | 5.4 | 2.9 | 4.1 | 7.2 | 3.2 | 1.1 | 0.8 |
| E484Q | 0.7 | 1.2 | 30.3 | 2.8 | >5840 | >1748 | >641 | 2.5 | 1.7 | 2.7 | 2.1 | 1.4 | 2.4 | 1.4 | 1.4 | 9.4 | 1.6 | 1.3 | 2.0 | 2.2 | 3.7 | 5.2 | 1.3 | 0.5 | 0.9 |
| F486V | 0.5 | 0.7 | 1.2 | 0.7 | >4213 | >1023 | 1.4 | 64.2 | >79 | 148.5 | 58.7 | 101.1 | 0.3 | 0.6 | 0.8 | 305.3 | 7.1 | 1.1 | 2.6 | 1.8 | 3.3 | 1.7 | 1.2 | 0.9 | 1.2 |
| F490S | 1.1 | 1.2 | 399.1 | 173.2 | >4213 | >691 | 8.5 | 1.4 | 1.3 | 1.1 | 1.3 | 0.7 | 2.4 | 0.7 | 0.7 | 4.3 | 1.1 | 1.2 | 1.0 | 0.5 | 2.3 | 2.6 | 1.4 | 1.2 | 1.0 |
| Q493K | 0.7 | 0.9 | 1.0 | 1.0 | >3422 | 4.9 | >466 | >1469 | 4.1 | 3.2 | 20.0 | 222.5 | 0.9 | 0.9 | 1.1 | >1871 | 6.7 | 1.0 | 1.6 | 2.0 | 2.6 | 1.3 | 1.8 | 1.3 | 0.9 |
| S494P | 1.2 | 1.2 | 7.9 | 0.7 | >4425 | 1.2 | >636 | >1238 | 0.5 | 0.8 | 0.7 | 1.4 | 0.9 | 1.4 | 1.0 | 13.3 | 0.7 | 1.0 | 0.9 | 1.4 | 1.3 | 0.9 | 0.8 | 1.1 | 1.0 |
| N501Y | 0.6 | 0.9 | 0.5 | 2.2 | 2.2 | 1.7 | 1.5 | 3.4 | 1.1 | 0.9 | 0.7 | 16.0 | 0.8 | 0.8 | 1.0 | 2.0 | 0.7 | 1.0 | 0.8 | 0.7 | 1.5 | 1.4 | 0.9 | 1.4 | 1.1 |
| P681H | 1.0 | 1.2 | 2.9 | 2.6 | 1.3 | 1.1 | 1.6 | 1.4 | 1.3 | 1.0 | 3.1 | 1.1 | 1.0 | 1.2 | 1.1 | 1.6 | 1.5 | 1.0 | 1.0 | 1.0 | 1.3 | 1.3 | 1.3 | 1.8 | 1.7 |
| D936Y | 1.0 | 0.9 | 0.7 | 0.5 | 0.9 | 1.0 | 0.6 | 0.6 | 0.7 | 0.7 | 0.4 | 0.7 | 0.8 | 0.9 | 0.7 | 1.1 | 0.7 | 1.2 | 0.8 | 0.9 | 1.1 | 1.4 | 0.9 | 1.1 | 1.0 |

**Figure S2. Heat map showing nAb potency as the ratio between the $IC_{50}$ values of pseudoviruses with single substitutions or deletions against $IC_{50}$ of WT pseudoviruses for each antibody.** Ratios are color coded as follows: <10, green = no major impact; 10-50 yellow = moderate loss of potency and >50 red = highly resistant. <0.2, blue = increased potency. Combinations not determined are shown in gray (n.d.) Data shown represent at least two independent experiments each with an intra-assay duplicate, except where indicated by *.
* indicates only tested once.

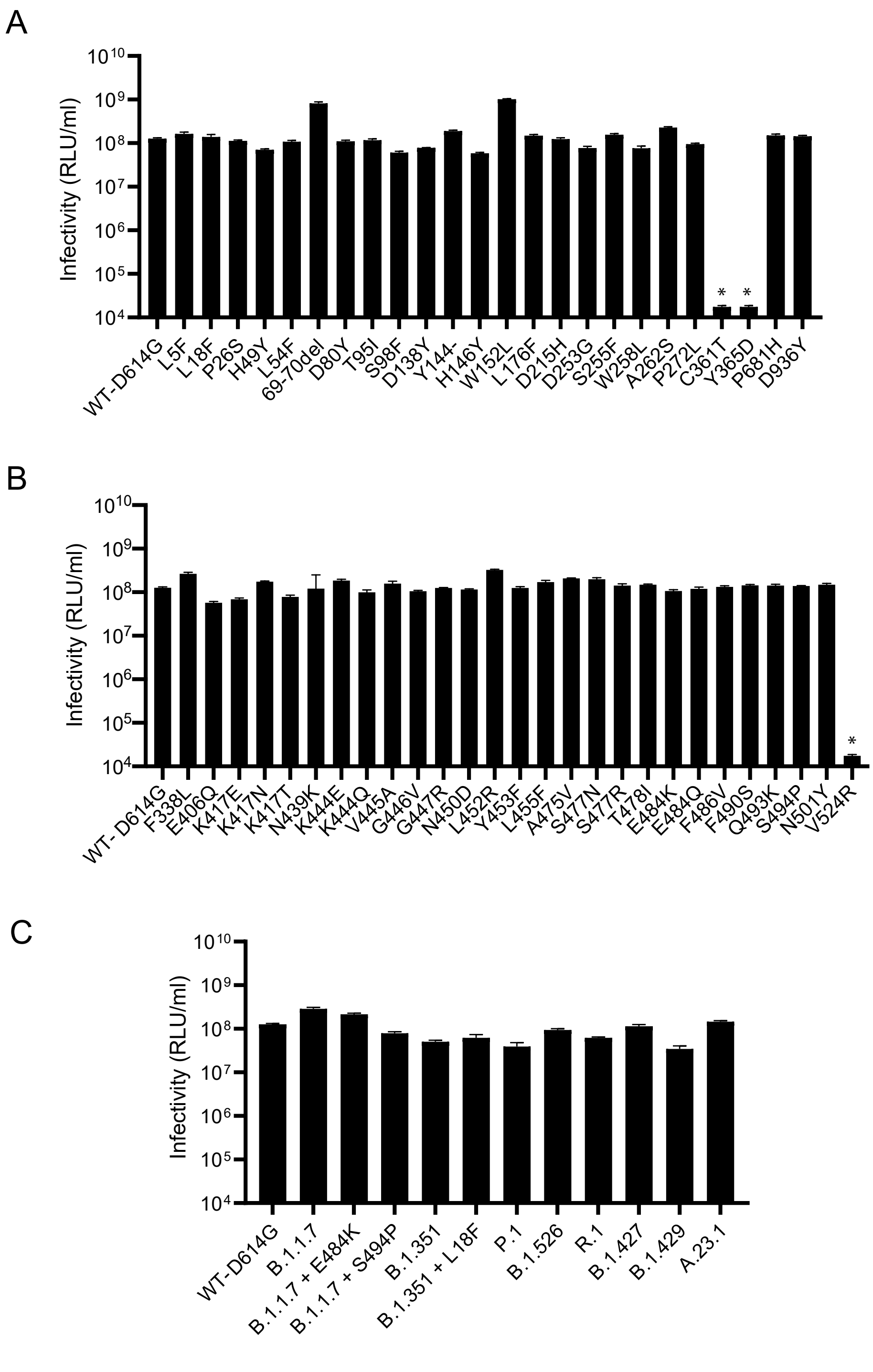


**Figure S3. Infectivity of the pseudoviruses bearing Spikes with the single and multiple amino acid substitutions**. (A) Infectivity of pseudoviruses with deletions (del) or single substitutions in the NTD and S2 regions of Spike. (B) Infectivity of pseudoviruses with single substitutions in the RBD. (C) Infectivity of pseudoviruses with multiple substitutions representing VOCs or VOIs. Volume-normalized transfection supernatants were used to inoculate stable 293T-ACE2/TMPRSS2 cells. Results shown are averages of two independent experiments. Infectivity expressed as relative luminescence units per ml (RLU/ml) with standard deviation. Data shown represent at least two independent experiments each with an intra-assay duplicate. * indicates pseudoviruses with low infectivity that could not be used for neutralization studies.

**Table S1. SARS-CoV-2 Variants**

| VARIANT NAME PANGOLIN [WHO LABEL] (NEXTSTRAIN) | LOCATION OF EARLIEST DOCUMENTED SAMPLES | SPIKE SUBSTITUTIONS |
|---|---|---|
| B.1 | Germany | D614G |
| B.1.1.7 [Alpha] (20I/501Y.1) | United Kingdom | 69-70deletion, 144deletion, **N501Y,** A570D, D614G, P681H, T716I, S982A, D1118H |
| B.1.1.351 [Beta] (20H/501Y.V2) | Republic of South Africa | (L18F)*, D80A, D215G, Del241-243, **K417N**, **E484K**, **N501Y**, D614G, A701V |
| P.1 [Gamma] (20J/501Y.V3) | Brazil/Japan | L18F, T20N, P26S, D138Y, R190S, **K417T**, **E484K**, **N501Y**, D614G, H655Y, T1027I, V1176F |
| B.1.427/B.1.429 [Epsilon] (20C/S:452R) | California | S13I, (P26S)*, W152C, **L452R**, D614G |
| B.1.526 [Iota] (20C/S:484K) | New York | L5F, T95I, D253G, **E484K**, D614G, A701V |
| A.23.1 (19B) | Uganda | F157L, **V367F,** Q613H, P681R |
| R.1 (20B) | Arizona | W152L, **E484K,** D614G, G769V |

* Pseudoviruses with or without the indicated Spike substitution shown in parenthesis were tested and showed no difference for our panel of therapeutic antibodies.
**Notes that the spike used in the pseudovirus contained D614G rather than D614.
Bold letters indicate substitutions in the receptor binding domain.
